# Optical nanoscopy reveals SARS-CoV-2-induced remodeling of human airway cells

**DOI:** 10.1101/2021.08.05.455126

**Authors:** Wilco Nijenhuis, Hugo G.J. Damstra, Emma J. van Grinsven, Malina K. Iwanski, Patrique Praest, Zahra E. Soltani, Mariëlle M.P. van Grinsven, Jesse E. Brunsveld, Theun de Kort, Lisa W. Rodenburg, Dorien C.M. de Jong, Henriette H.M. Raeven, Sacha Spelier, Gimano D. Amatngalim, Anna Akhmanova, Monique Nijhuis, Robert Jan Lebbink, Jeffrey M. Beekman, Lukas C. Kapitein

## Abstract

A better understanding of host cell remodeling by the coronavirus SARS-CoV-2 is urgently needed to understand viral pathogenesis and guide drug development. Expression profiling and electron microscopy have frequently been used to study virus-host interactions, but these techniques do not readily enable spatial, sub-cellular and molecular analysis of specific cellular compartments. Here, we use diffraction-unlimited fluorescence microscopy to analyze how SARS-CoV-2 infection exploits and repurposes the subcellular architecture of primary human airway cells. Using STED nanoscopy, we detect viral entry factors along the motile cilia of ciliated cells and visualize key aspects of the viral life cycle. Using Tenfold Robust Expansion (TREx) microscopy, we analyze the extensively remodeled three-dimensional ultrastructure of SARS-CoV-2-infected ciliated cells and uncover Golgi fragmentation, emergence of large and atypical multivesicular bodies enclosing viral proteins, ciliary clustering, and remodeling of the apical surface. These results demonstrate a broadly applicable strategy to study how viruses reorganize host cells with spatial and molecular specificity and provide new insights into SARS-CoV-2 infection in primary human cell models.

## Introduction

COVID-19 is an respiratory illness caused by infection with severe acute respiratory syndrome coronavirus 2 (SARS-CoV-2) [1–5]. Since its emergence in late 2019, the SARS-CoV-2 outbreak has developed into a global pandemic with severe medical and socioeconomic consequences [6–8]. Despite the rapid development of several effective vaccines, pharmacotherapeutic interventions to treat COVID-19 are urgently required [9]. It is therefore crucial to develop a better understanding of key virus-host interactions, such as entry, replication, intracellular trafficking and egress, in a clinically relevant model system.

As a β-coronavirus, SARS-CoV-2 comprises an enveloped single-stranded RNA. The 30-kb viral genome is decorated by the nucleocapsid (N) protein, whereas the viral membrane is decorated by the structural proteins envelope (E), membrane (M) and spike (S) [10–12]. Viral entry is mediated by the binding of spike to the human host receptor angiotensin converting enzyme II (ACE2) and subsequent proteolytic activation of spike by host proteases, including TMPRSS2 [4, 13]. Two-thirds of the viral genome consists of two overlapping ORFs encoding the polyproteins pp1a and pp1b, which are proteolytically cleaved to give rise to 16 non-structural proteins (nsps) that mediate various functions, including genome replication, minus-strand RNA synthesis and the production of subgenomic RNAs [12].

Earlier electron microscopy (EM) studies investigating the replication of other coronaviruses revealed extensive ultrastructural rearrangements involving multiple organelles including the endoplasmatic reticulum (ER) and the Golgi apparatus [10, 14, 15]. This has led to detailed insights into the architecture of the viral replication organelle (RO), where the viral RNA synthesizing machinery associates with ER membranes. Viral RNA synthesis is contained within double membrane vesicles, which may function to shield the viral RNA from the innate immune receptors that are activated by the double stranded RNA (dsRNA) intermediates of viral RNA replication [16, 17]. A molecular pore provides an exit from the double membrane vesicles for viral RNA, allowing replicated genomes to be translated to form new nucleocapsids and subgenomic RNAs [18]. Viral membrane proteins are then synthesized on the ER, incorporated into transport vesicles at ER exit sites and trafficked to the endoplasmic-reticulum-Golgi intermediate compartment (ERGIC) [19], where virus budding is believed to take place [10, 20, 21]. Whereas the architecture of the replication organelles has been extensively studied, many other aspects of the SARS-CoV-2 viral life cycle, including viral entry, budding and egress, are still unknown. In particular, it remains unclear how the trafficking of virus progeny affects cellular organization.

Ultrastructural analyses of viral infection are hampered by the disruption of recognizable morphological features in late stages of viral infection, which precludes identification based on morphology and necessitates immunolabeling of specific proteins to obtain a molecular understanding. In addition, reliable identification of viral particles based on morphology has proven problematic [22–25]. Immuno-EM can be used to localize specific proteins at the ultrastructural level, but is technically challenging and suffers from low labeling efficiency, a small field of view and a lack of volumetric information [26]. These limitations can be partially overcome by combining the large field of view and specific labeling offered by optical immunofluorescent microscopy with EM-based ultrastructural analyses, as was recently used to reveal a lysosomal, Arl8b-dependent exocytic pathway for the egress of SARS-CoV-2 [27]. However, such studies are labor-intensive and are limited by the resolution of conventional optical microscopy. To better understand virus-host interactions, combining ultrastructural information with super-resolution fluorescence imaging of specific proteins would be required, preferably in a physiologically relevant model system.

Here, we set out to visualize key aspects of the SARS-CoV-2 life cycle in primary human airway epithelium with nanoscopic resolution. Using STimulated Emission Depletion (STED) microscopy, we studied the spatial organization of viral structural proteins, non-structural proteins and dsRNA at <100 nm resolution. In addition, for a comprehensive understanding of the morphological changes of airway epithelium induced by SARS-CoV-2 infection, we used a novel type of Expansion Microscopy (Tenfold Robust Expansion (TREx) Microscopy [28]) in combination with both selective and non-selective protein stains [28, 29]. This enabled the volumetric visualization of epithelial ultrastructure with specific labeling of viral proteins and revealed extensive intracellular reorganization, including Golgi fragmentation, the emergence of large multivesicular bodies containing spike-positive vesicles (MVBs), clustering of cilia, and apical membrane remodeling. As such, this work reveals the power of volumetric fluorescence nanoscopy for studying SARS-CoV-2-induced cytopathic effects in primary cell culture models.

## Results

### Nanoscopic visualization of the SARS-CoV-2 infection cycle in Vero E6 cells

To validate our infection model, we first infected Vero E6 cells with two independent SARS-CoV-2 strains (clinical isolates NL/2020 and Utrecht p1). Both strains productively infected the culture, as demonstrated by a positive staining for the viral spike protein in the majority of cells at 1 day post infection (dpi) at a multiplicity of infection (MOI) of 0.01 (Figure S1). We then proceeded to visualize key aspects of the viral infection cycle by co-staining infected Vero E6 cells for different viral and organelle markers. To distinguish between early and late stages of infection, we co-stained for spike, dsRNA and the *cis*-Golgi marker GM130, as disruption of the Golgi apparatus is indicative of late-stage coronavirus infection [30, 31]. Uninfected cells were devoid of dsRNA puncta and displayed normal Golgi organization (Figure 1A inset 1). Cells at early stages of infection could be identified by low numbers of ROs, slight dispersion of the Golgi and decoration of the Golgi by low levels of spike (Figure 1A inset 2). Cells at late stages of infection contained large numbers of perinuclear ROs, were intensely stained by spike and showed a severely disrupted Golgi (Figure 1A inset 3).

**FIGURE 1:**
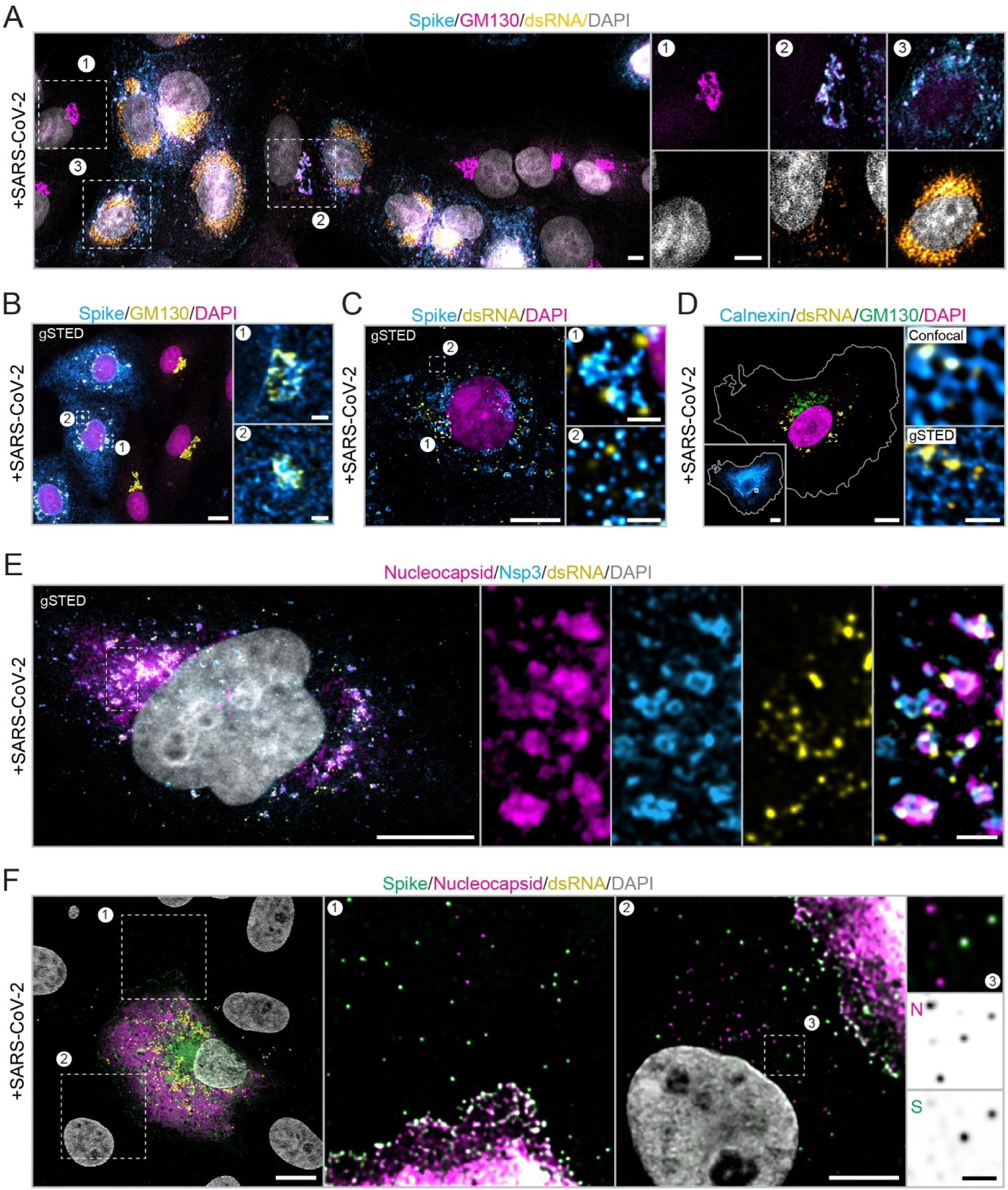
Nanoscopic visualization of the SARS-CoV-2 infection cycle in Vero E6 cells. (A-F) Immunofluorescent imaging of indicated markers in SARS-CoV-2 infected Vero E6 cells showing (A) progressive increase of viral markers and disruption of Golgi morphology (GM130), (B) disruption of Golgi in later stages of infection (Figure shows 3 infected cells next to 3 uninfected cells), (C) association of spike-positive vesicles with ROs (dsRNA), (D) association of ROs with ER (Calnexin) in early stages of infection, (E) intricate morphology of viral replication machinery (nsp3), ROs and nucleocapsid, and (F) viral shedding. Scale bars are 10 µm (overviews A-F), 5 µm (panels F1, F2) or 1 µm (panels F3 and zooms in A-E). MOI: 0.01. Cells were fixed at 1 dpi or 2 dpi (B).

To better examine changes in cellular architecture upon infection, we turned to super-resolution microscopy (Figure 1B). Using STED microscopy, dispersed Golgi fragments often appeared round and swollen and some fragments were intertwined with other spike-positive structures, suggesting that spike accumulates in Golgi-associated structures other than the *cis*-Golgi at later stages of infection. Indeed, in late stages of infection spike localized to multiple other compartments (Figure 1B,C) and was often associated with ROs marked by dsRNA (Figure 1C inset 1). ROs originate from the ER and are surrounded by remodeled, ‘zippered’ ER in the form of convoluted membranes [16]. To visualize the association of ROs with the ER, we stained for dsRNA and the luminal ER marker calnexin. Subsequent STED microscopy revealed that ROs were closely associated with calnexin-positive ER tubules, but not surrounded by calnexin (Figure 1D), suggesting that the ER-derived membranes known to enclose ROs might be devoid of typical ER-associated proteins.

Earlier work on other coronavirus has used immuno-EM to map specific markers of the viral assembly pathway to specific elements within the RO ultrastructure [14–16]. This revealed that dsRNA exclusively localizes within double membrane vesicles, nsp3 accumulates on convoluted membrane, and nucleocapsid localizes to both convoluted membrane and associated double membrane spherules. To map these structural elements for SARS-CoV-2, we visualized these markers using multi-color STED microscopy (Figure 1E). Consistent with the reports mentioned above, we found that dsRNA, nsp3 and nucleocapsid localize to separate, but closely associated structures. Remarkably, nsp3 was organized in polygon-shaped structures, which were diffusely labeled by nucleocapsid protein and associated with multiple ROs (Figure 1E). The size and shape of nsp3 polygons was reminiscent of the organization of the ER tubules proximal to ROs (Figure 1D), suggesting that nsp3-positive structures are shaped by the ER, as proposed previously [16].

Finally, to visualize viral shedding and entry, we stained for the structural viral proteins spike and nucleocapsid in combination with dsRNA and examined the immediate periphery of isolated highly infected cells (Figure 1F). Often, cells adjacent to highly infected were devoid of dsRNA puncta, indicating that viral replication had not yet begun, but contained many small spike- and nucleocapsid-positive puncta with a highly uniform size and intensity, consistent with individual viral particles (Figure 1F, insets 1 and 2). Whereas all puncta stained positively for nucleocapsid, some puncta appeared to have selectively lost spike (Figure 1F, inset 3). This suggests that these spike-negative puncta are uncoated, internalized virions at the earliest stage of infection. Thus, by imaging specific organelle and viral markers (Golgi morphology, number of ROs and distribution of spike) with STED microscopy, we could distinguish between levels of infection, characterize the localization of spike and dsRNA and their respective association with the Golgi and the ER at a nanoscopic level, map structural elements of the SARS-CoV-2 RO and detect viral shedding and entry.

### Characterization of SARS-CoV-2 infection and subcellular localization of viral entry factors in a human airway model

We next examined SARS-CoV-2 infection of primary human airway cells because the upper respiratory system is the primary site of droplet-based infection by airborne viruses. Previously, we established a robust protocol to differentiate human nasal epithelial cells (HNECs) from nose brushes of healthy donors (Figure 2A) [32]. As ciliated cells are the main permissive cell type for SARS-CoV-2 in the respiratory epithelium [4, 33–42], we chose a differentiation protocol that enriched for ciliated cells. We validated the composition of our cultures by staining with cell type specific markers (Figure S2) and found a mix of basal cells (∼20-30% of total), MUC5AC^+^ goblet cells (∼10-20%), ciliated cells (∼40-50%) and sparse CC10^+^ club-like cells (Figure 2B).

**FIGURE 2:**
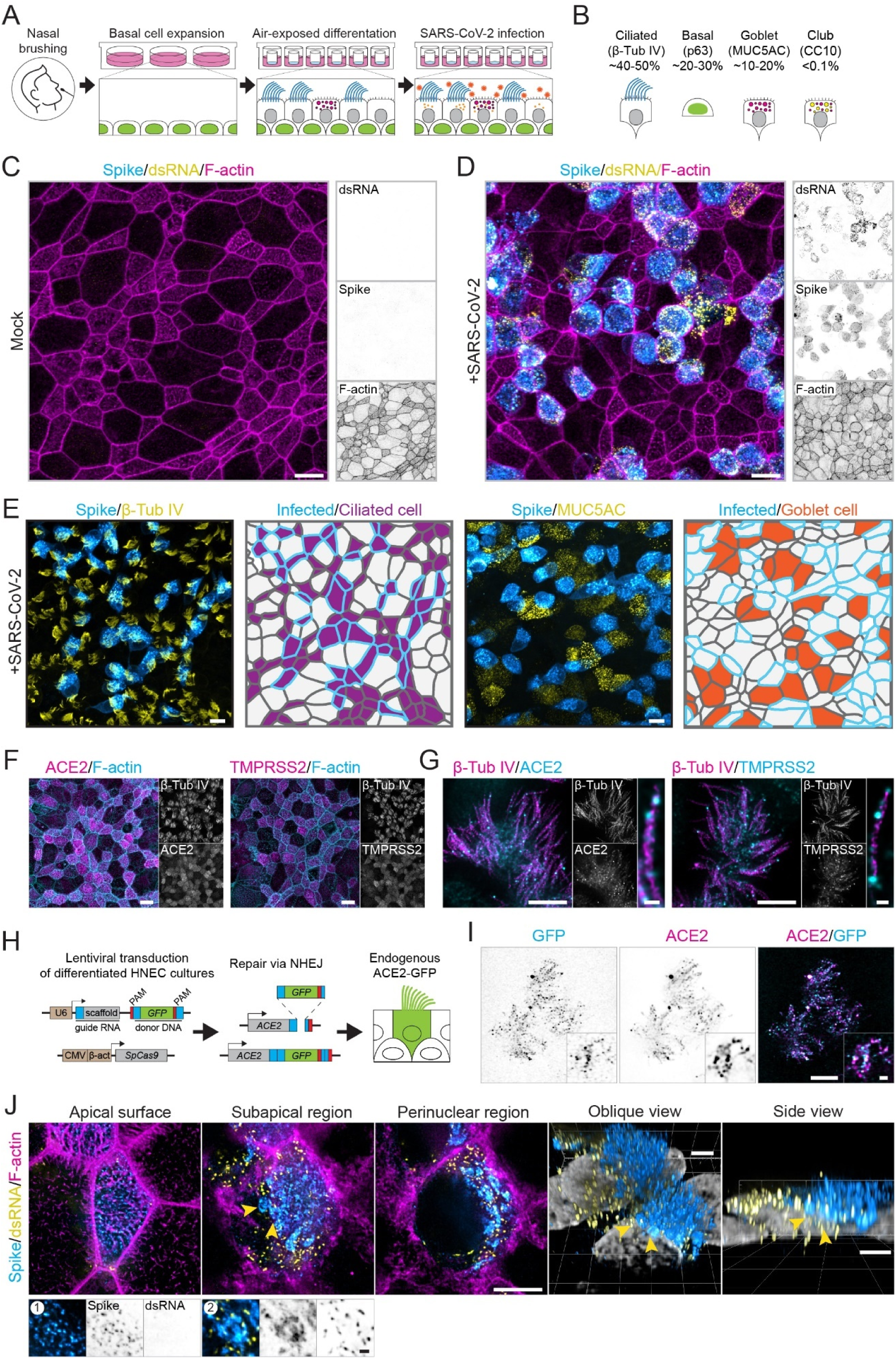
Preferential infection of ciliated cells in a human airway model. Overview (A) and characterization (B) of HNEC culture system and SARS-CoV-2 infection model. (C,D) Immunofluorescent confocal imaging of uninfected (C) and SARS-CoV-2 infected (D) HNECs stained for indicated viral markers and F-actin (Phalloidin). (E) Immunofluorescent imaging and segmentation of infected HNECs stained for Spike and cell type markers β-Tub-IV (ciliated cells) or MUC5AC (goblet cells), showing preferential infection of ciliated cells. (F,G) Immunofluorescent confocal (F) and gSTED (G) imaging of F-actin, cilia and viral entry factors ACE2 and TMPRSS2 in HNECs showing enrichment of entry factors on cilia. (H-I) Strategy for (H) and immunofluorescent imaging of (I) endogenous GFP-tagging of ACE2 in HNECs using ORANGE-based genome editing, showing that immunolabeling of ACE2 and GFP labels overlapping punctate structures along cilia (see also G). (J) Immunofluorescent gSTED imaging and renders of SARS-CoV-2 infected HNECs, showing Spike-positive puncta at the apical surface and enrichment of closely associated replication organelles (dsRNA) and Spike-positive organelles in the subapical and perinuclear regions. Insets show details from single optical sections from the apical surface (J1) and the subapical region (J2). Scale bars are 10 µm (C-E, F), 5 µm (G, I, J), 3 µm (renders) or 0.5 µm (zooms G, I, J). MOI: 0.1. Cells were fixed at 3 dpi.

Fully differentiated HNEC cultures were inoculated apically with SARS-CoV-2 from one of two clinical isolates (NL/2020 or Utrecht patient 1) at an MOI of 0.2 for 2h, before washing and incubating for up to 3 dpi before fixation. At 3 dpi, patches of infected cells could be found among regions devoid of infected cells (Figure 2C and 2D). By correlating spike expression with cell type-specific markers, we found that only ciliated cells were infected (Figure 2E). Consistently, ciliated cells are the primary target of SARS-CoV-2 in most reported airway models [4, 33–42] although infection of goblet cells [4, 33, 34, 38, 43], club cells [4, 37, 39, 41, 44] and basal cells [40, 41, 45] has also been observed, particularly as secondary targets in later stages of infection [41]. This could imply that our airway model system mostly resembles early stages of infection.

Since we observed preferential infection of ciliated cells, we wondered whether the SARS-CoV-2 host entry factors ACE2 and TMPRSS2 were also enriched in ciliated cells. Whereas ACE2 has been reported to enrich apically on ciliated cells [39, 45–48] and to decorate cilia [47], the localization of TMPRSS2 has remained unclear. We observed that both ACE2 and TMPRSS2 were enriched at the apical surface of ciliated cells (Figure 2F). Moreover, both entry factors were also enriched on the cilia themselves and formed bright puncta along their length (Figure 2G). To confirm this distribution, we endogenously tagged the cytosolic tail of ACE2 with GFP in HNECs using ORANGE-based genome editing [49] (Figure 2H) and found that ACE2-GFP puncta colocalized with direct antibody labeling of the ACE2 ectodomain (Figure 2I).

We then explored the distribution of spike and dsRNA within ciliated cells with STED microscopy (Figure 2J). Spike was found in various patterns: as puncta at the apical surface, as larger structures and as clusters of puncta associated with ROs in the subapical and perinuclear regions, consistent with our observations in Vero E6 cells. ROs were enriched in the subapical and perinuclear regions and mostly absent from the apical surface. Whereas STED microscopy provided a comprehensive overview of the spatial distribution of viral markers in flat adherent cells such as Vero E6 cells, visualizing the distribution of viral markers in 3D was more challenging in thick airway cells due to the diffraction-limited axial resolution of conventional 2D gSTED microscopy. This was especially apparent when rendering the cellular volume from the side (Figure 2J), showing the axially elongated point-spread function. Therefore, a different strategy was needed to study SARS-CoV-2 infection with nanoscopic resolution in complex three-dimensional cell types.

### TREx reveals the 3D ultrastructure of human airway cells

In recent years, expansion microscopy (ExM) has emerged as a technique that enables nanoscale imaging of biological samples with conventional microscopes. In ExM, specimens are embedded in a swellable polymer gel to physically expand the sample and isotropically increase resolution in *x*, *y* and *z* [50]. The resolution increase is partly limited by the expansion factor of the gel, which is 4.5x for the original ExM protocol [51]. Recently, we co-developed TREx microscopy: a variation on the original ExM protocol that provides tenfold expansion and can be readily combined with small molecule stains to directly label lipid membranes or provide a total protein stain, thereby offering ultrastructural context (Figure 3A) [28]. With TREx, we set out to expand HNEC cultures directly from Transwell filters to visualize cellular morphology in 3D.

**FIGURE 3:**
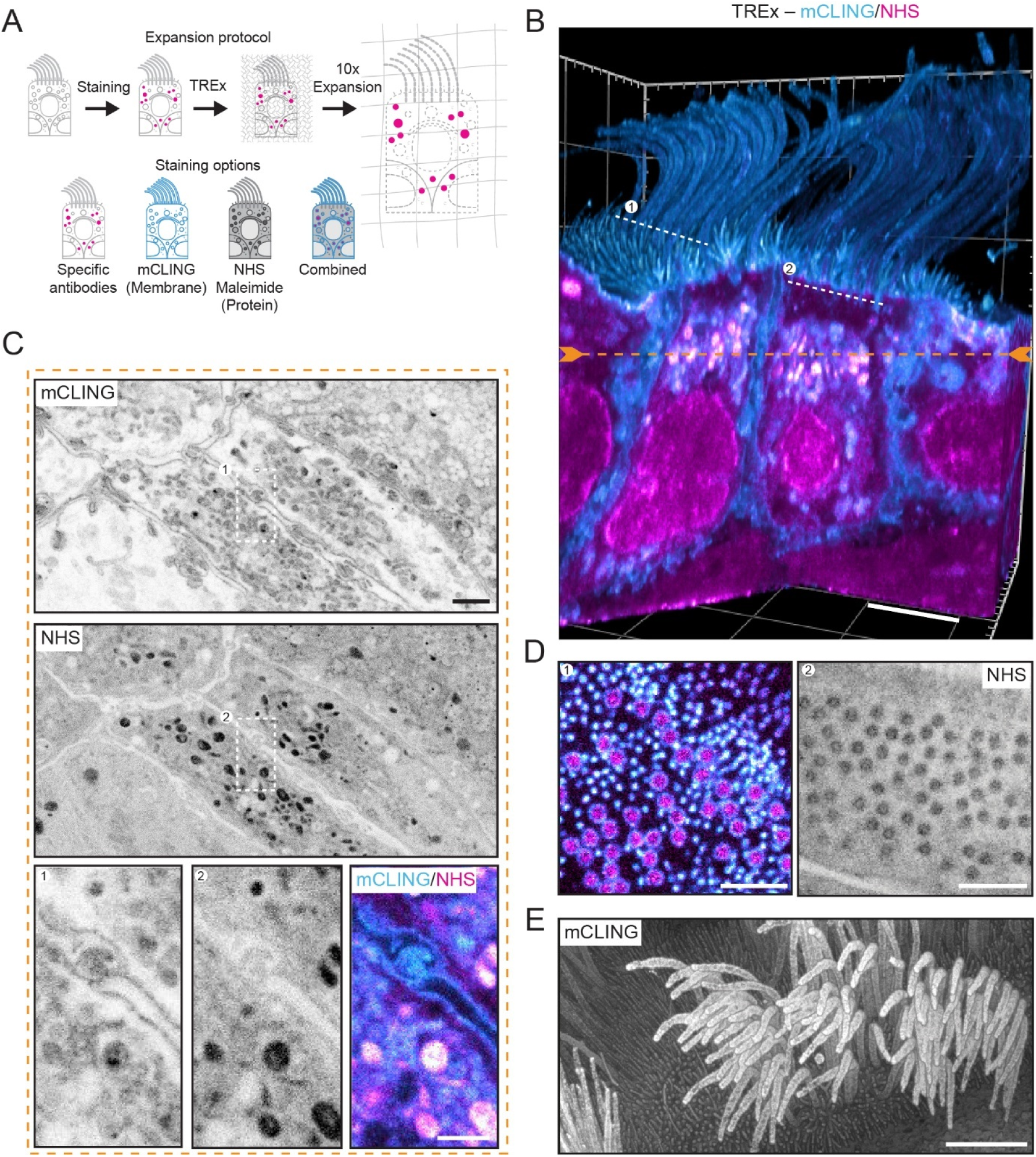
TREx microscopy reveals ultrastructure of human airway cells. (A) Ten-fold Robust Expansion (TREx) Microscopy principle. HNECs are stained with specific antibodies or small molecule stains (mCling, NHS, maleimide) and embedded in a swellable polymer gel that isotropically expands ten-fold after proteolytic digestion. (B-E) Multiple renders of TREx of HNECs stained with mCLING (total membrane) and NHS-ester (total protein), highlighting different aspects of cellular morphology for the same cellular volume. See also movie S1. (B) Volumetrically rendered side-view, clipped to reveal intracellular ultrastructure. Dotted orange and white lines mark planes shown in C and D, respectively. (C) Single plane from B. Zoom of representative region shows elaborate interdigitated membrane contacts between cells. (D) Single planes from (B) showing basal bodies. (E) Top-down depth-encoded volumetric render of (B) providing contrast reminiscent of scanning electron microscopy. Scale bars (corrected to indicate pre-expansion dimensions): B, E ∼2 μm, C, D ∼1 μm, zooms ∼500 nm.

To label cellular membranes, we stained HNECs with mCLING, a palmitoylated small peptide that intercalates into the lipid bilayer and is compatible with TREx chemistry [28, 52], and combined this with a fluorophore-labeled N-hydroxysuccinimide (NHS) ester or maleimide that covalently binds to lysines or cysteines, respectively, and thereby provides density-based ultrastructural context [29]. Using glutaraldehyde-based fixation optimized to preserve membrane morphology, followed by gel polymerization inside the Transwell chamber and subsequent expansion, we visualized motile cilia (captured mid-beat) extending from the apical surface (Figure 3B and movie S1). Optical sectioning revealed a variety of structures, including extensive interdigitated membrane contacts between adjacent cells (Figure 3C). Furthermore, mCLING staining of cilia was hollow with the ciliary lumen labeled by the total protein stain, allowing for the easy identification of intracellular and extracellular compartments (Figure 3D). In addition, specific structures within the total protein stain could be identified based on the density of staining, such as the basal bodies at the base of the cilia (Figure 3D). Finally, by rendering the same dataset as a depth-coded volumetric projection we could obtain a detailed overview of apical morphology (Figure 3E), resembling scanning electron microscopy. Thus, TREx enables volumetric imaging of the nanoscale organization of membranes or proteins within complex cells and tissue and facilitates flexible representation of the data, either volumetric or as single sections.

### Super-resolution imaging reveals the emergence of enlarged CD63-positive multivesicular bodies in SARS-CoV-2 infected human airway cells

Because infection disrupted the organelle distribution and morphology in Vero E6 cells, we next examined intracellular reorganization in infected HNECs. First, we characterized Golgi morphology in HNEC cultures by gSTED microscopy and TREx microscopy. Whereas in non-infected cells the Golgi was localized in perinuclear ribbon-like structures, infected cells contained small Golgi fragments that were dispersed throughout the cytoplasm (Figure 4A). These fragments appeared intertwined with clusters of spike-positive densities (Figure 4A, B and movie S2), which could be visualized in 3D using TREx microscopy. To further explore the intracellular organization of infected HNECs, we used a general protein stain in combination with the selective labeling of spike and GM130 (Figure 4C). Again, we observed clusters of spike-positive densities often in contact with the Golgi, but the general protein stain did not provide additional structural information about these structures (Figure 4C, inset 1). This total protein stain did, however, reveal a different type of compartment in infected cells that was not observed in uninfected cells (Figure 4C-E and movie S3). These protein-dense, large, round organelles with heterogenous content were devoid of GM130 staining and typically contained various sparse spike densities.

**FIGURE 4:**
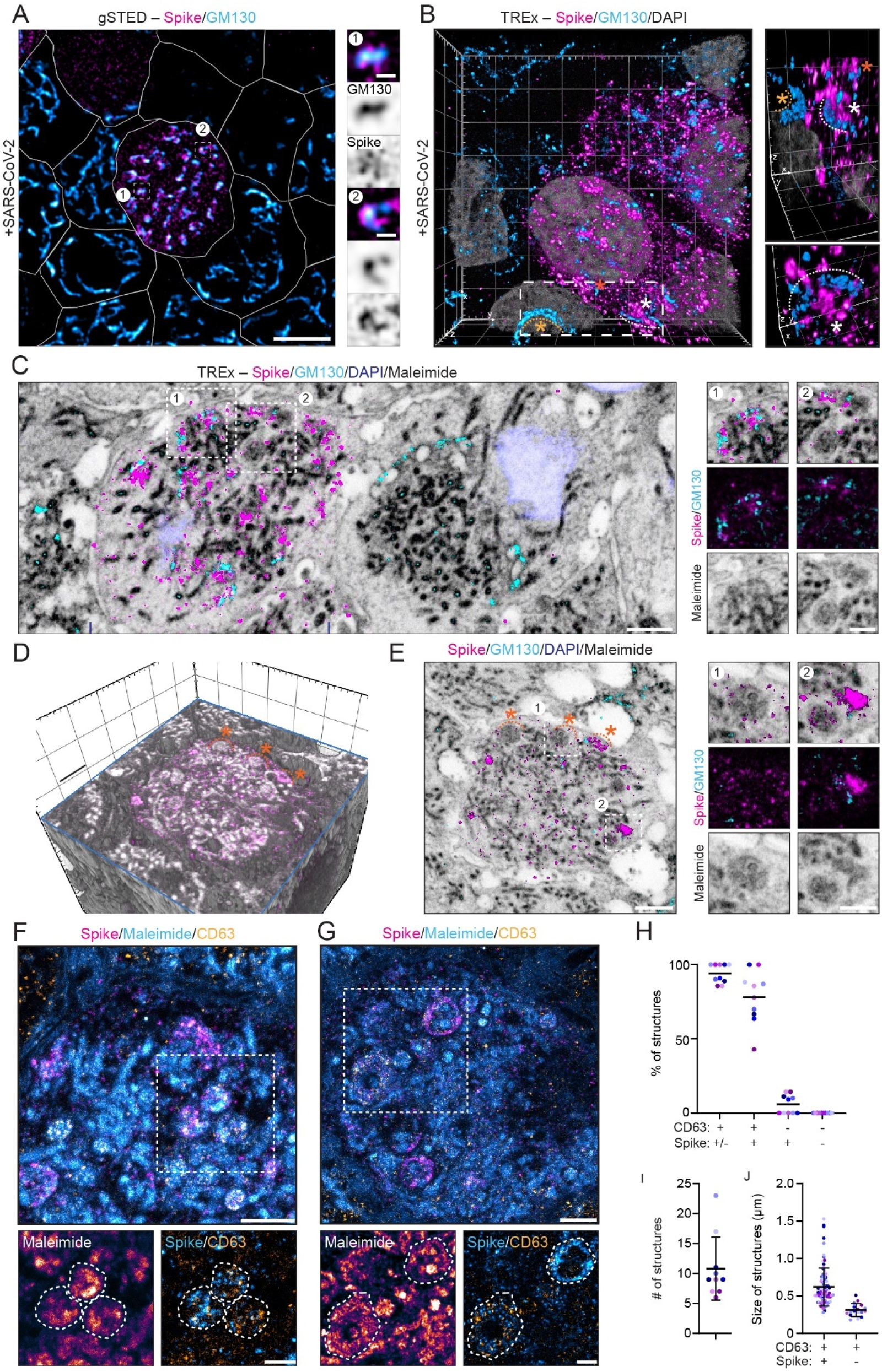
TREx reveals intracellular reorganization in SARS-CoV-2 infected human airway cells. (A, B) gSTED (A) and TREx (B) imaging of spike and Golgi (GM130) in SARS-CoV-2 infected ciliated cells, showing close association of spike with Golgi and dispersal of the Golgi apparatus in infected cells. See also movie S2. (C-E) TREx imaging of Spike, Golgi, nuclei (DAPI) and total protein (maleimide) in SARS-CoV-2 infected ciliated cells. (C) Top-down view of a section revealing ultrastructural reorganizations in an infected cell. Representative zooms 1 and 2 show dispersed Golgi apparatus (1) and large, round, densely-stained virus-induced organelles with irregular morphology (2). (D and E) TREx imaging of an infected cell shown as (D) a 3D volumetric render, computationally unroofed to show ultrastructural rearrangements and as (E) a single plane. Zooms show spike-positive and spike-negative maleimide densities, resembling organelles shown in zoom C2. Asterisks indicate morphologically similar reorganizations. See also movie S3. (F and G) TREx imaging as in (E), stained for spike and intralumenal vesicles (CD63) with zooms showing accumulation of spike in late endosomal compartments containing CD63-positive intralumenal vesicles (F) and peripheral enrichment of CD63 and spike in swollen late endosomal compartments (G). (H-J) Quantification of large virus-induced organelles for 108 organelles in 10 cells, 2 independent replicates. Scale bars are 5 µm (A), ∼1 μm (B-G), 500 nm (all zooms). TREx scale bars were corrected to indicate pre-expansion dimensions. MOI: 0.1. Cells were fixed at 3 dpi.

To elucidate the identity of these structures, we reasoned that the heterogenous content could be intralumenal vesicles. Consistently, enlarged endolysosomal compartments positive for the intralumenal vesicle marker CD63 have been reported in cells infected with the α-coronavirus porcine epidemic diarrhea virus (PEDV) [53]. We therefore visualized CD63 together with spike and the total protein stain in HNEC using TREx microscopy (Figure 4 F,G). Careful examination of the ultrastructure revealed that on average 94±6% of the SARS-CoV-2-induced structures were positive for CD63, 78±18% were positive for both CD63 and spike and 7±7 % were only positive for spike, whereas no structures were observed that were labeled by neither CD63 nor spike (n = 108 structures from 10 cells (2 independent replicates) (Figure 4 H,I). CD63 and spike were typically detected throughout these large compartments (Figure 4F) but were sometimes enriched at the periphery of the largest of these structures (Figure 4G). Size measurements of all detected structures in infected cells furthermore revealed that structures containing both CD63 and spike were on average 2 times larger than structures with only CD63 (Figure 4 I,J). These results indicate that SARS-CoV-2 infection results in the formation of expanded multivesicular bodies (MVBs), a type of late endosomes.

We hypothesized that the formation of these large compartments must be reflected in a general reorganization of late endosomes throughout the cellular volume. To visualize this more comprehensively and better compare CD63 distribution in infected and uninfected cells, we stained CD63 in infected Vero E6 cells (Figure 5A). In uninfected cells, CD63 localized to dispersed punctate organelles which were enriched in the perinuclear region, as is typical for MVBs. In infected cells, most CD63 signal was instead found on large spherical organelles. As in the airway cells, these swollen compartments contained various spike densities, but were not specifically enriched in spike. The compartments were also not directly associated with ROs (Figure 5A). Whereas CD63 staining could be found within these compartments, it was often also enriched in the periphery of the organelle (Figure 5A), which is atypical for MVBs.

**FIGURE 5:**
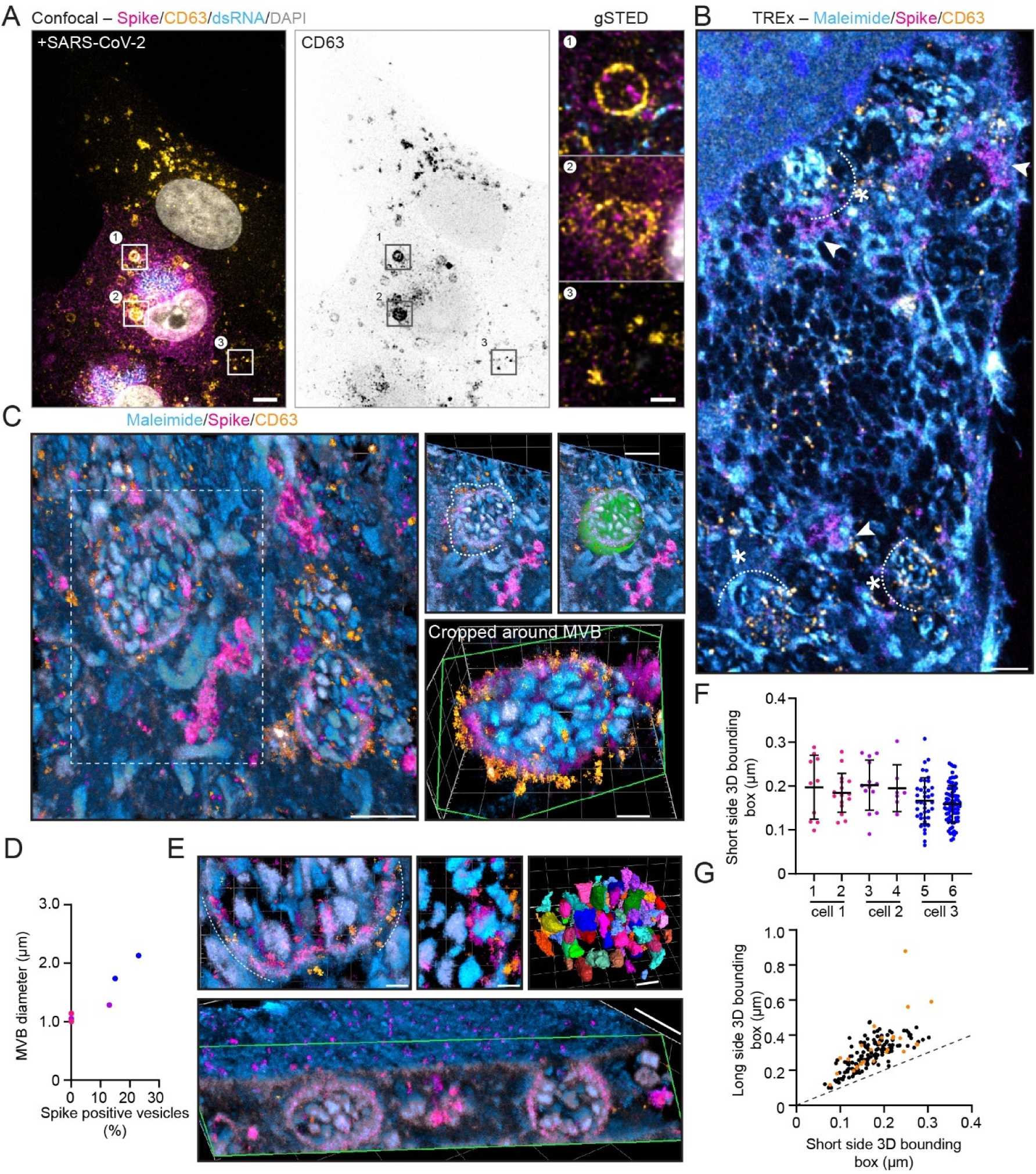
Nanoscopic characterization of SARS-CoV-2-induced atypical multivesicular bodies. (A) gSTED imaging of spike, dsRNA and intralumenal vesicles (CD63) in Vero E6 cells, showing swelling of multivesicular bodies in SARS-CoV-2 infected cells (insets 1 and 2). (B) TREx imaging of spike and CD63 of samples as in (A), showing swollen compartments containing CD63-positive intralumenal vesicles (asterisks) and other spike-containing densities (arrows). For full field of view see Figure S3B. (C) Volumetrically rendered and computationally unroofed detail from TREx imaging as in (B), showing a large swollen multivesicular bodies (MVBs) similar to the example in inset A1, morphologically segmented along the limiting membrane (middle and right panel above, green) and shown in isolation (lower panel) with a perpendicular clipping plane (green). See also movie S4. (D) Quantification of MVB diameter at the widest point (corrected to indicate pre-expansion dimensions) versus the percentage of spike-positive intralumenal vesicles (ILV) inside the MVB, color coded per cell (n = 6 MVBs from 3 cells). (E) Top row: Zoom of ILVs near the rim of the MVB, detail of Spike positive ILV, and visualization of segmented ILVs (see methods). Bottom: Side view of dataset shown in (C) showing one of the MVBs continuous with the plasma membrane. (F) Quantification of the short side of the 3D bounding box of each segmented ILV (see E) separated per MVB (color-coded as in D, n = 10, 16, 13, 7, 41, 78 ILVs for MVB 1-6, respectively, from 3 cells, corrected to indicate approximate pre-expansion dimensions). (G) length of long versus short side of the 3D bounding boxes of pooled ILVs from (F), dashed line indicates spherical particles, Spike positive ILVs are indicated in orange. Scale bars are 5 µm (A), 1 µm (Zooms in A, B, C, below side view E), 300 nm (zoom in C and right top panel E), 200 nm (left top and middle zooms in E). TREx scale bars were corrected to indicate pre-expansion dimensions. MOI: 0.01. Cells were fixed at 2 dpi.

When we used TREx microscopy to image CD63 and spike in combination with the intracellular ultrastructure, we found that our general protein stain clearly labeled vesicle-like intralumenal material in the enlarged CD63-positive organelles (Figure 5B,C and S3A,B). These enlarged, vesicle-filled organelles were strikingly similar to structures seen in recently published EM images of cells infected with SARS-CoV-2 or mouse hepatitis virus, which were interpreted as lysosomes filled with viral particles based on the presence of the late endosome/lysosome marker LAMP1 and the morphology of the intralumenal material (see Figure 2 of [27]). However, LAMP1 also labels MVBs and intraluminal vesicles can be easily mistaken for viral particles [22–25], necessitating a better identification of the intralumenal material through the selective labeling of spike. This revealed that the percentage of spike-positive vesicles per CD63-positive MVB varied between 0 and 23%, with larger MVBs having a larger percentage (Figure 5D, E). Remarkably, spike decorated the outside of these intralumenal structures, suggesting that these structures might represent individual viral particles en route to either lysosomes or the plasma membrane (Figure 5E). Indeed, of the two MVBs shown in Figure 5E, one appears continuous with the plasma membrane, consistent with release of intralumenal content (see also Video 4).

Unlike conventional intralumenal vesicles, which are mostly spherical, but similar to the particles described in late endosome-like structures (see Figure 2 of [27]), the intralumenal content in the virus-induced MVBs was shaped irregularly (Figure 5E-G). Our three-dimensional data set enabled us to quantify the shape by measuring the long and short sides of the bounding boxes enclosing these structures (Figure 5F,G). This revealed that most particles were ellipsoidal with a long side of 300±10 nm and a short side of 170±5 (average± s.d., n= 165 particles from 6 MVBs of 3 cells). Together, these results demonstrate the importance of augmenting morphological observations with specific labeling and reveal that SARS-CoV2 infection leads to the formation of large and atypical CD63-positive MVBs, in which a subset of intraluminal particles were spike-positive vesicles.

### Super-resolution imaging reveals ciliary disorganization in SARS-CoV-2 infected human airway cells

Cohesive ciliary beating is essential for the clearance of mucus from the airway and ciliary defects are a typical consequence of respiratory infection [42, 54]. Furthermore, we observed that the viral entry factors were enriched on cilia (Figure 2F,H). We therefore next examined ciliary morphology upon SARS-CoV-2 infection. We first visualized ciliary morphology by staining β-IV-tubulin and imaged these samples using gSTED microscopy. As a result of the fixation protocol and sample mounting for STED imaging, cilia were flattened and spread out across the apical surface. In uninfected samples, this helped us visualize clearly separated, healthy cilia (Figure 6A). Upon SARS-CoV-2 infection however, cilia appeared clustered (Figure 6B) and individual cilia could often not be resolved. Infection-induced ciliary loss or shortening, which has been reported in some cases [4, 34, 42], was not readily apparent. Interestingly, the ciliary clustering defect was also observed in uninfected cells within the same region. We hypothesized that ciliary clustering could be caused by extracellular virions that crosslink cilia, but found that spike was not enriched on cilia and instead localized to the apical membrane of infected cells (Figure 6C). To further examine ciliary morphology, we visualized the full apical surface in 3D using TREx imaging of total membranes (Figure 6D,E). Cilia in uninfected samples remained upright and curved uniformly, as expected for beating cilia. This was in stark contrast with infected samples, where cilia were clearly clustered at the tips, irregularly shaped and disorganized.

**FIGURE 6:**
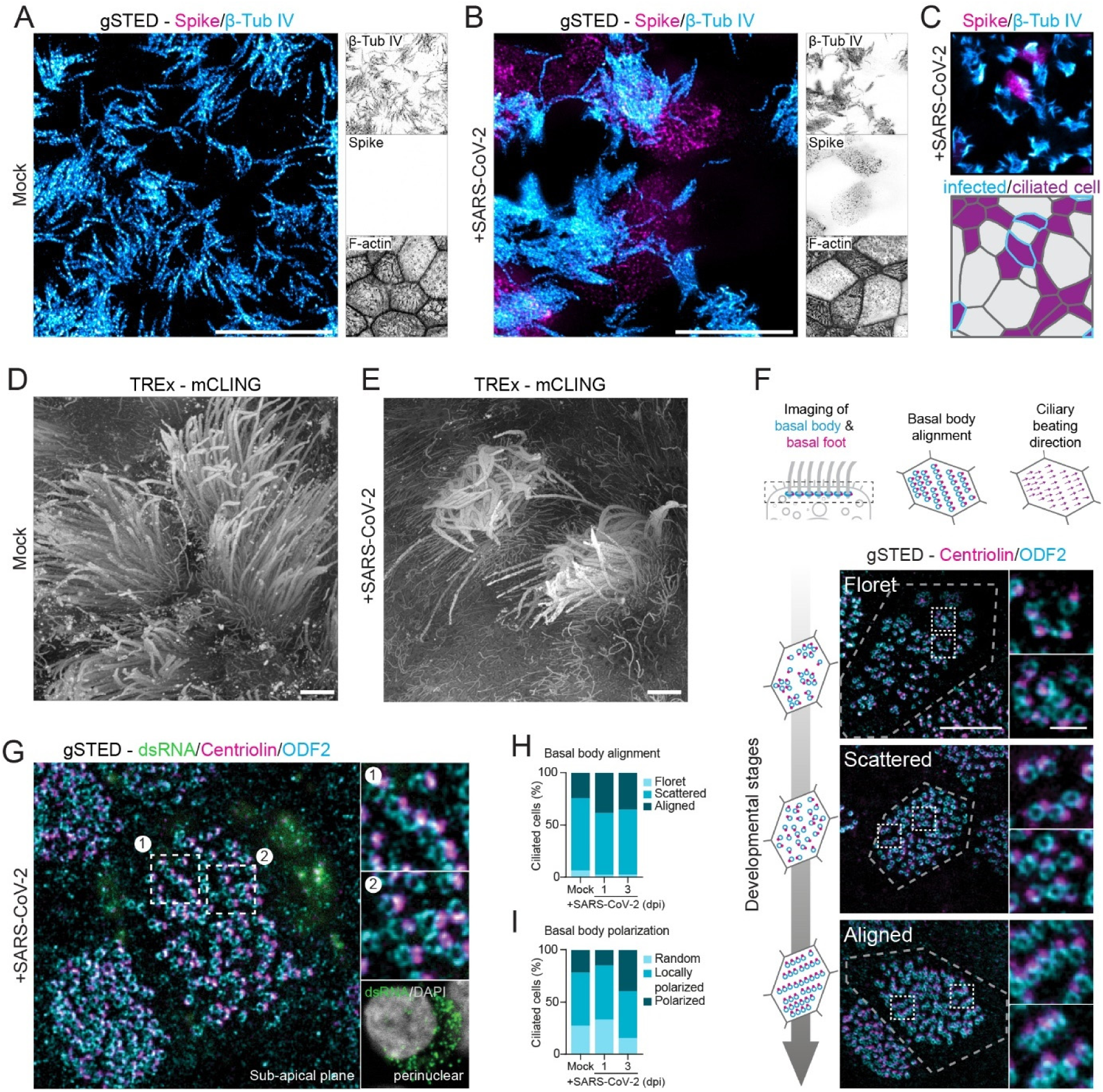
Super-resolution imaging reveals ciliary clustering in SARS-CoV-2 infected human airway cells. (A,B) gSTED imaging of Spike, cilia (β-Tub-IV) and F-actin in mock (A) and SARS-CoV-2 infected (B) HNECs showing ciliary clustering in infected samples. (C) Confocal imaging and segmentation of samples as in (B) showing ciliary clustering phenotype in spike negative cells. (D, E) Depth-coded volumetric render of representative mock (D) and SARS-CoV-2 infected HNECs (E). (F-H) Analysis of basal body alignment and ciliary beating direction on ciliated HNECs by gSTED imaging of ODF2 (basal bodies) and centriolin (basal foot). (F) Development stages of basal body organization. (G) gSTED imaging of centriolin and ODF2 in dsRNA positive, SARS-CoV-2-infected HNECs with no apparent changes in either basal body alignment or polarization. (H, I), Quantification of basal body alignment (H), derived from ODF2 distribution, and basal body polarization (I), derived from ODF2-centriolin axis alignment, in infected and uninfected HNECs. Scale bars are 10 µm (A-C), 5 µm (F, G overviews), ∼2 µm (D, E, corrected to indicate pre-expansion dimensions), 1 µm (F, G zooms). or 0.5 µm (zooms G, I, J). MOI: 0.1. Unless indicated otherwise, cells were fixed at 3 dpi.

To achieve coordinated ciliary beating, cilia positioning is organized through the positioning of basal bodies, centriole-like structures at the base of each cilium [55]. To test if ciliary clustering was related to basal body reorganization, we analyzed basal body organization in mock-infected and SARS-CoV-2-infected cells. During development, ciliated cells undergo three stages of basal body arrangement – floret, scattered and aligned – to achieve a uniform beating direction [55]. These stages are distinguished by the alignment of the basal bodies in combination with the orientation of the basal foot relative to the basal body, which indicates the ciliary beating direction (Figure 6F). To analyze basal body alignment and polarization, we stained for ODF2 and centriolin, markers for the basal body and basal foot, respectively (Figure 6G-I). As ciliary clustering was observed throughout the infected culture, independent of spike signal (Figure 6C), we included all ciliated cells in infected HNEC cultures in our analysis. This analysis revealed that infection did not affect basal body alignment or polarization, even in highly infected cells (Figure 6G). Thus, SARS-CoV-2 infection induces ciliary clustering and disorganization, but this is not a result of basal body disorganization.

### Nanoscopic visualization of surface remodeling of SARS-CoV-2 infected cells

While imaging infected Vero E6 cells with TREx, we noted that infected cells were rounded up and decorated by a large number of filopodia-like membrane protrusions and occasional larger membrane sheets (Figure 7A). Spike was distributed across the surface of infected cells, but was enriched on the tips of these protrusions. In contrast, spike only sparsely decorated the surface of neighboring cells without cytosolic spike, suggesting that the strong spike labeling on infected cells resulted from viral egress and shedding, rather than entry. This is consistent with our observation of spike at the limiting membrane of MVBs (Figure 5E) and the observed fusion of MVBs with the plasma membrane. Furthermore, recent work has showed similar actin-rich protrusions decorated with viral particles and implicated these in egress or cell-to-cell spreading [56]. To further examine surface reorganization in infected Vero E6 cells, we next performed gSTED microscopy of actin and found that uninfected Vero E6 cells have lower levels of filamentous actin and a smoother cortex than infected cells (Figure S4). Infection induced many more actin bundles and more intense stress fibers, as well as an increase in filopodia-like protrusions (Figure S4).

**FIGURE 7:**
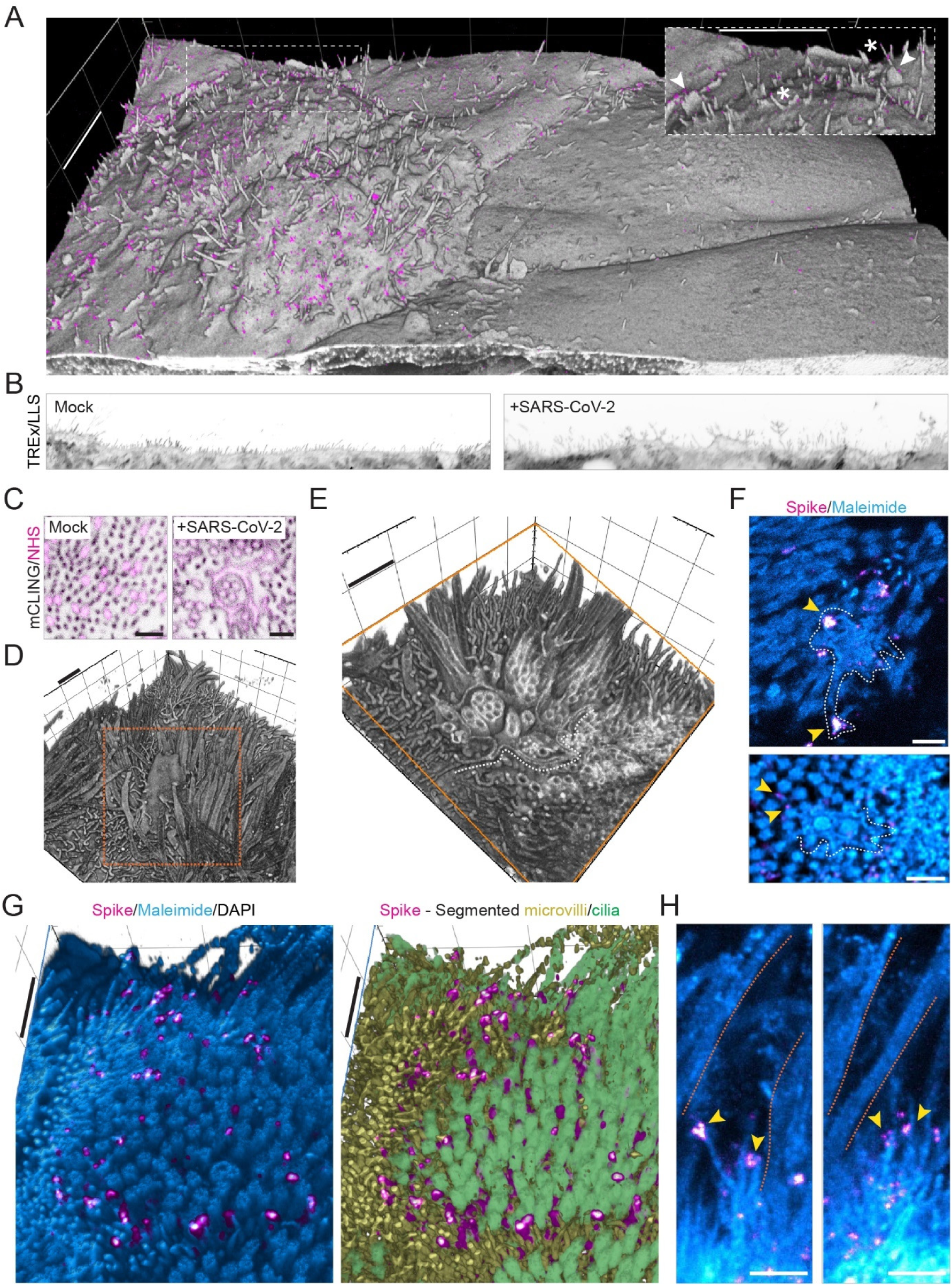
SARS-CoV-2 induces spike-positive protrusions. (A) Depth-coded volumetric render of TREx imaging of a group of infected (spike-positive) and uninfected (spike-negative) Vero E6 cells, showing association of spike with virus-induced filopodia. Insert is zoom of boxed region, arrows point to membrane sheets, asterisks indicate spike positive protrusions. See also movie S5. (B) TREx/Lattice-light sheet imaging of apical surface in mock and SARS-CoV-2 infected HNECs, showing tree-like apical protrusions in the infected sample. (C) Single plane of a representative apical region of mock or SARS-CoV-2 infected ciliated cells showing extensive membrane (mCLING) remodeling surrounding multiple cilia. (D and E) TREx 3D volumetric render (D) and unroofed zoom (E) of SARS-CoV-2 infected HNEC sample showing an extreme example of a remodeled membrane engulfing multiple cilia. Clipping plane shown in orange. White dotted line marks partial outline of the membrane sack. (F) TREx imaging of spike and total protein (maleimide) in infected ciliated cells showing similar reorganizations as in C and D. White lines trace the membrane, yellow arrows indicate spike puncta. (G) TREx imaging of an infected mCLING-labeled ciliated cell clipped perpendicular to cilia to show spike puncta colocalizing on apical membrane protrusions. Right: same view segmented to differentiate between microvilli and cilia. (H) Representative single planes of spike-positive apical protrusions. Orange lines mark cilia, yellow arrows point to Spike puncta. See also movie S6. Scale bars are 4 µm (A), ∼1 μm (B, D, F), 500 nm (C, E, G). TREx scale bar was corrected to indicate pre-expansion dimensions. MOI: 0.01 (A), 0.1 (F-H), 0.2 (B-E). Cells were fixed at 1 dpi (A) or 3 dpi (B-H).

As cilia are interspersed with actin-rich microvilli on airway cells and we previously found that spike is enriched on the apical surface, we hypothesized that the apical surface of airway cells might also be remodeled upon infection. To visualize the effect of SARS-CoV-2 on the apical membrane topology of airway cells, we stained membranes with mCLING, followed by TREx microscopy. Lattice-light sheet imaging of these samples revealed that infection induced long, branched apical membrane structures (Figure 7B and S4). Occasionally, we also observed large sheet-like membrane extensions between and around motile cilia, consistent with the membrane sheets we observed on infected Vero E6 cells (Figure 7A,B). In extreme cases, this led to the formation of large, hollow membrane structures around cilia (Figure 7C) that extended to form membrane sacks encapsulating multiple cilia, thereby generating a multi-layered membrane structure (Figure 7D,E).

Direct visualization of the membrane using mCLING requires fixation with glutaraldehyde, which is incompatible with antibody-labeling of spike. This precluded us from determining the level of infection of these cells. To validate that the membrane sacks were induced by infection, we combined the spike staining with a fluorescent protein stain to visualize general morphology. In these samples, we observed similar apical membrane structures and found spike labeling colocalizing with the membrane sacks (Figure 7F). Moreover, the combination of spike and total protein stain in expanded samples allowed us to more closely examine the apical enrichment of spike that we had observed using gSTED (Figure 2J). Whereas gSTED imaging showed that spike abundantly decorated the apical region, the increased resolution of TREx revealed that near the apical surface, spike exclusively the decorated microvilli (Figure 7G). These findings were further supported by morphological segmentation of the apical total protein signal into microvilli and cilia (Figure 7G). Finally, by volumetric rendering, we were able to localize spike puncta along individual microvilli and occasionally found them clustered at the tip of branched microvilli (Figure 7G). These results indicate that, by reorganizing the actin cytoskeleton, SARS-CoV-2 infection induces large scale apical membrane remodeling in ciliated cells, including the emergence of spike-positive branched protrusions.

## Discussion

Here we have used innovative light microscopy to comprehensively visualize key aspects of the SARS-CoV-2 infectious cycle in Vero E6 cells and primary human airway cells. We applied recent advances in super-resolution fluorescence microscopy to image specific viral markers and target structures with <100-nanometer resolution in whole cells. Using multicolor immunofluorescent STED microscopy, we visualized viral uncoating during entry, the subdomain architecture of replication organelles and the distribution of host entry factors on cilia. In addition, the use of general protein and membrane labels in TREx microscopy enabled us to resolve cellular ultrastructure in complex 3D samples. This allowed us to bridge scales between visualizing large morphological defects, such as ciliary clustering, and altered suborganellar structures, such as spike-positive intralumenal vesicles. Moreover, by combining these general labels with selective stainings, we could map specific proteins within the cellular ultrastructure. This revealed Golgi fragmentation, emergence of large atypical MVBs and viral egress resulting in remodeled microvilli. Our work in human airway cells complements a recent study that used fourfold ExM to explore the RO in Vero E6 cells [57]. We expect that our methodology will be a valuable addition to existing techniques for visualizing viral infection in general and help further our understanding of SARS-CoV-2 induced cytopathy.

Our TREx protocol yields a single-step isotropic tenfold resolution improvement, which enables volumetric multicolor super-resolution imaging of thick human-derived samples on a conventional confocal microscope. This facilitates volumetric optical imaging of the 3D ultrastructure of human airway cells, in combination with markers for specific proteins. This allowed us to visualize specific viral markers in whole cellular volumes, which remains challenging with electron microscopy, despite recent advances such as correlative light and electron microscopy and 3D electron tomography [58, 59]. For example, we could identify the large organelles emerging upon infection as atypical CD63-positive MVBs and directly determine the percentage of spike-positive intralumenal vesicles. Importantly, recent work has used morphological assignment to report the emergence of virus-filled lysosomes upon SARS-CoV-2 infection [27]. Given the striking resemblance of these reported structures with what we have here identified as atypical CD63-positive MVBs, our results strongly suggest that the structures reported earlier as lysosomes are in fact large, atypical MVBs in which virions are at most a subset of the intralumenal particles. As such, our work demonstrates the importance of combining (three-dimensional) ultrastructural imaging with selective labeling.

Enlarged late endosomal compartments were previously observed using EM [53, 60], but have remained underappreciated in the absence of a more comprehensive overview of MVB morphology throughout infected cells. Whereas some viruses use MVBs to exit cells [61, 62], recent work reported that viral egress was unaffected by treatments aimed at blocking MVB fusion with the plasma membrane [27]. Nonetheless, we here observed the fusion of an MVB containing virus-like intraluminal particles with the plasma membrane, suggesting this could be a pathway for egress in unperturbed conditions. Alternatively, MVBs could mostly be an intermediate for viral trafficking between the Golgi/ERGIC and lysosomes, which have also been shown to mediate egress [27]. In addition, many spike-positive particles might not be complete virions, since recent work has shown that overexpressed spike can also end up within MVBs [63]. Future experiments are needed to explore the exact interplay between these different compartments during virion formation and egress.

In addition to intracellular reorganizations, we also observed severe morphological rearrangements at the cell surface. In Vero E6 cells, we observed overall surface roughening and the emergence of spike-positive filopodia. Smooth Vero E6 cells without filopodia were almost devoid of spike labeling both on the surface and in the cytoplasm. This indicates that these virus-induced filopodia result from viral replication or egress and are not related to viral entry. These findings are consistent with recent reports that SARS-CoV-2-infected Vero E6 and Caco-2 cells form an increased number of actin-rich filopodial protrusions that are branched and elongated compared to those on uninfected cells. These protrusions were decorated by viral particles and were postulated to be involved with viral egress [56, 64]. Indeed, other viruses are known to induce actin extensions, such as elongated microvilli or filopodia, for efficient cell-to-cell spread of virions [65]. Importantly, we recapitulated these findings in human ciliated airway cells, where we observed severe apical remodeling and spike accumulation on actin-rich microvilli. In these infected HNECs, we also observed clustering of ciliary tips in both infected and neighboring uninfected cells. This was not caused by basal body disruption nor by direct crosslinking of cilia by viral particles, as we did not observe a clear accumulation of spike on cilia. Ciliary clustering could be caused by a more global antiviral response, potentially related to interferon signaling. Elucidating how innate immune responses prevent the spread of viral infection in airway cultures will be an important direction for future work.

We studied infection in human nasal epithelial cultures because the upper airway is the primary target of SARS-CoV-2 [66]. Organotypic cultures have become a critical tool in drug development and pre-clinical and cell biological research, and are increasingly being adopted for use in virological research [67]. Through recent work using human airway models, it has become clear that viral tropism, the degree of infection and the replication efficiency depend on the airway model used (nasal, tracheal or bronchial), differences in culture protocols, donor-to-donor variation and the specific SARS-CoV-2 strain used for infection [68]. These factors together affect the efficiency of viral infection and the severity of cytopathic effects. Because our culture protocol allows for the robust growth and differentiation of basal cells obtained from nose brushes of healthy donors and patients suffering from COVID-19, the nanoscopy approaches developed here could aid the comparison of HNECs derived from asymptomatic and severely affected COVID-19 patients, which may provide new insights into COVID-19 progression and could help guide the development of therapeutics. In addition, these imaging tools will be widely useful for the study of human-derived tissue models.

## Material and Methods

### Key resources table

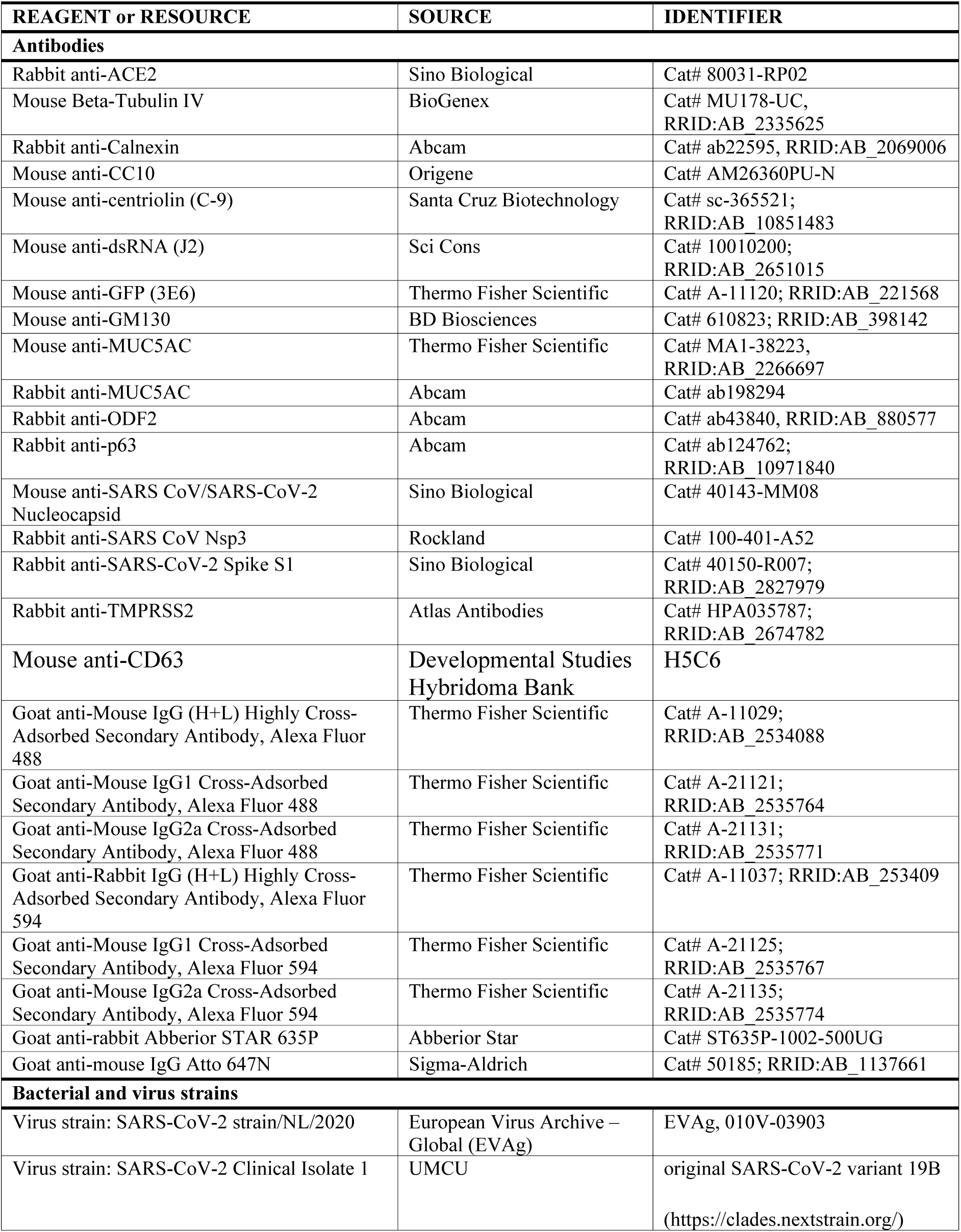
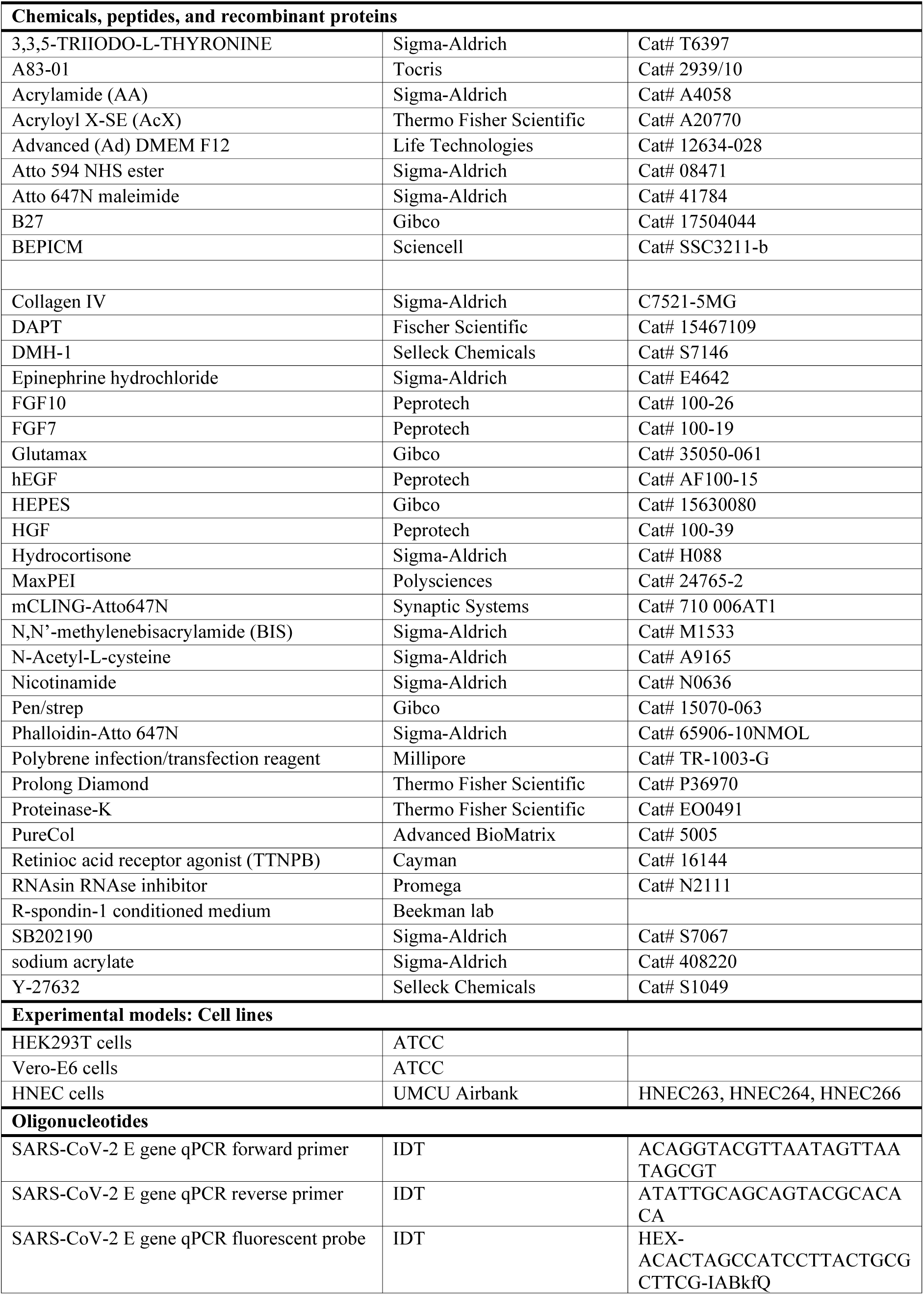
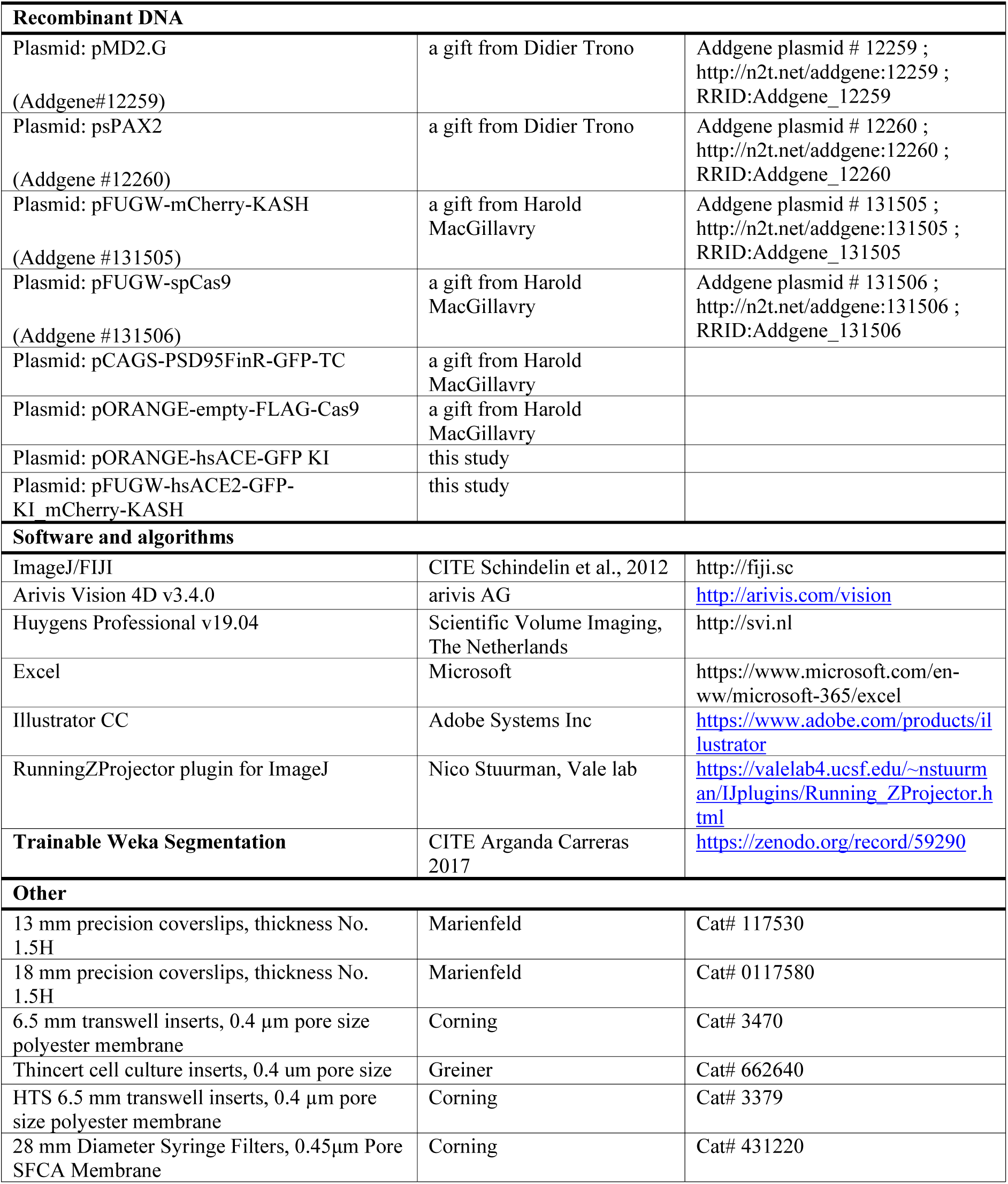

### Cells

Vero E6 cells and HEK293T cells were maintained in Dulbecco’s modified Eagle’s medium (DMEM, Invitrogen) supplemented with 10% fetal calf serum (FCS, PAA laboratories), 100 Units/ml penicillin, 100 µg/ml streptomycin and 2mM L-glutamine.

### Human airway epithelial cells culture and differentiation

Adult human airway cells were collected from nasal brushings of healthy volunteers without respiratory tract symptoms. All donors provided explicit informed consent and the study was approved by the Medical Research Ethics Committee (Tc-BIO protocols 16-856 and 20/806). The material used in this study was derived from two independent donors. All data was derived from donor HNEC0266 with the exception of data shown in figure 3, which was derived from donor HNEC0263. Human nasal epithelial cells (HNEC) were cultured as described previously [32]. In brief, HNEC were expanded on collagen IV-coated (50 ug/ml, Sigma Aldrich)-6-well culture plates using basal cell (BC) expansion medium consisting of BEPICM (50%, Sciencell #3211), Advanced (Ad) DMEM F12 (23,5%, Life Technologies #12634-028), R-spondin-1 conditioned medium (20%), HEPES (10 mM, Gibco #15630080), Glutamax (1x, Gibco #35050-061), Pen/strep (1x, Gibco #15070-063), B27 (2%, Gibco #12587010), Hydrocortisone (0.5 mg/mL, Sigma #H088), 3,3,5-triiodo-L-thyronine (100 nM, Sigma #T6397), Epinephrine hydrochloride (0.5 mg/mL, Sigma #E4642), N-Acetyl-L-cysteine (1,25 mM, Sigma #A9165), Nicotinamide (5 mM, Sigma #N0636), A83-01 (1 mM, Tocris #2939/10), DMH-1 (1 uM, Sellech Chemicals, #S7146), Y-27632 (5 uM, Selleck Chemicals #S1049), SB202190 (500 nM, Sigma #S7067), HGF (25 ng/mL, Peprotech # 100-39), hEGF (5 ng/mL, Peprotech #AF100-39), DAPT (Fischer Scientific #15467109), FGF7 (25 ng/mL, Peprotech #100-19) and FGF10 (100 ng/mL, Peprotech #100-26). To generate air-liquid-interface (ALI) 2D cultures, cells were dissociated with TrypLE express enzyme and seeded onto PureCol-coated (30 mg/mL, Advanced BioMatrix)-6.5 mm transwell inserts (0.4 um pore size polyester membrane, Corning #3470) and cultured in submerged conditions in BC expansion medium. When confluent, cells were differentiated in ALI-differentiation medium consisting of A83-01 (50 nM, Tocris #2939), hEGF (0.5 ng/mL, Peprotech #AF10015), 3,3,5-Triiodo-L-Thyronine (100 nM, Sigma #T6397), Epinephrine hydrochloride (0.5 mg/mL, Sigma #E4642), Retinioc acid receptor agonist (100 nM, Cayman #16144), Hydrocortisone (0.5 mg/mL, Sigma #H088) and Pen/Strep (1%, Gibco #15070-063) in 492,5 mL AdDMEM/F12 supplemented with additional A83-01 (500 nM, Tocris #2939). After 3-4 days of complete submerged culture conditions, the volume of medium at the apical side was decreased to a minimal amount for cells to be submerged. After 4 days, ALI-differentiation medium was removed and refreshed with ALI-differentiation medium supplemented with DAPT (5 mM, Fisher Scientific #15467109) for 14-18 days. The addition of DAPT to the differentiation medium together with the small amount of medium supplemented to the apical side of the culture shifted the differentiation program towards ciliated cells. Medium was refreshed twice a week (Mondays and Fridays), at which point the apical side was washed with PBS to remove mucus.

### Viruses

SARS-CoV-2 strain/NL/2020 (EVAg, 010V-03903) and SARS-CoV-2 Clinical Isolate 1 were fully sequenced (original SARS-CoV-2 variant 19B (https://clades.nextstrain.org/) and cultivated on monkey kidney Vero E6 cells. All work with SARS-CoV-2 was conducted in Biosafety Level-3 conditions at the University Medical Center Utrecht, according to WHO guidelines (Laboratory biosafety guidance related to coronavirus disease (COVID-19)). The respective virus titer was determined by titration of infectious particles on Vero E6 cells (TCID50) and quantitative real-time reverse transcription-PCR (qRT-PCR) specific for the SARS-CoV-2 E gene [69].

**Table 1:**
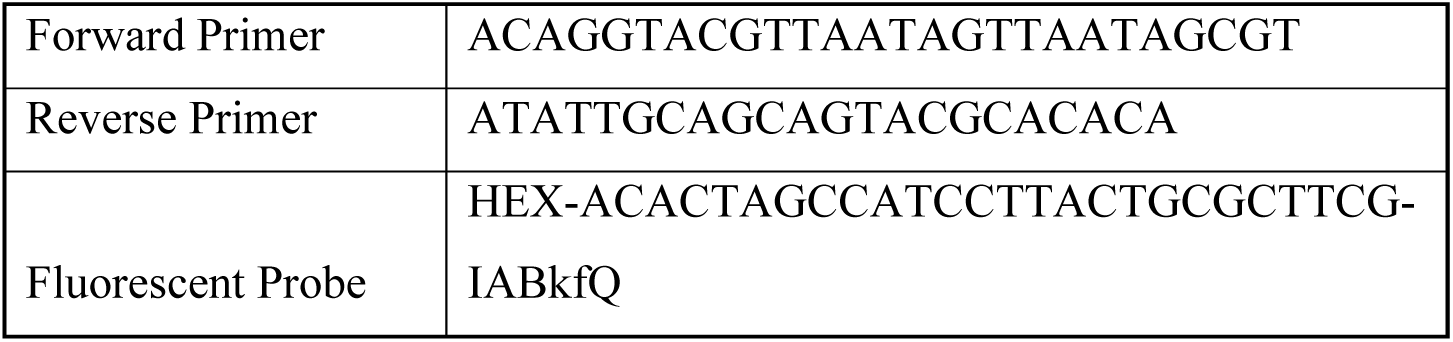
Primer and Probe sequences SARS-CoV-2 E gene

### Infections

Vero E6 were seeded the day prior to infection on 18 mm precision coverslips (thickness No. 1.5H, #0117580, Marienfeld) and maintained DMEM supplied with 2% FCS, 100 Units/ml penicillin, 100 µg/ml streptomycin and 2mM L-glutamine. The viruses were inoculated at the indicated multiplicity of infection (MOI) for 2h at 37°C. After inoculation the virus was removed, the cells were rinsed with PBS and fresh medium was applied.

Differentiated HNECs were incubated with phosphate-buffered saline (PBS) for 20 minutes at 37°C to remove the mucus from the apical surface before inoculation with the respective viruses. The viruses were diluted in advanced DMEM/F-12 and inoculated apically at the indicated MOI for 2h at 37°C. After inoculation, the virus was removed and the apical side was carefully rinsed with PBS three times, followed by PBS removal and apical exposure to air.

### Plasmids and cloning

pMD2.G (Addgene#12259) and psPAX2 (Addgene #12260) were gifts from Didier Trono. pFUGW-mCherry-KASH (Addgene #131505,[49]) and pFUGW-spCas9 (Addgene #131506, [49]), pCAGS-PSD95FinR-GFP-TC and pORANGE-empty-FLAG-Cas9 were gifts from Harold MacGillavry. pORANGE-hsACE-GFP KI, a plasmid for C-terminal endogenous tagging of ACE2 using the ORANGE method of CRISPR/Cas9-based genome editing [49], was generated by ligating a primer dimer encoding the target sequence TCTGAACATCATCAGTGTTT into the BbsI sites of pORANGE-empty-FLAG-Cas9. Subsequently, this plasmid was digested and a GFP donor module designed to ligate between codons 796 and 797 of ACE2 and flanked by TCTGAACATCATCAGTGTTTTGG sequences, encoding the target sequence and protospacer adjacent motif (PAM), was generated by PCR from pCAGS-PSD95FinR-GFP-TC and ligated into the HindIII and XhoI sites. pFUGW-hsACE2-GFP-KI_mCherry-KASH, a lentiviral vector for C-terminal endogenous tagging of ACE2 was generated by subcloning the target sequence, gRNA scaffold and EGFP donor sequence, surrounded by target sequences, from pORANGE-hsACE-GFP KI into the PacI site of pFUGW-mCherry-KASH, by Gibson assembly cloning.

### Lentiviral infection and gene editing

Lentivirus packaging was performed by using MaxPEI-based co-transfection of HEK293T cells with psPAX2, pMD2.G and the lentiviral vectors pFUGW-hsACE2-GFP-KI_mCherry-KASH and pFUGW-spCas9. Supernatant of packaging cells was harvested up to 72 h after transfection, filtered through a 0.45-µm filter and incubated with a polyethylene glycol (PEG)-6000-based precipitation solution overnight at 4°C. After precipitation, virus was concentrated up to 100× by centrifugation and dissolution in 1× PBS. Before infection, differentiated HNEC cultures were incubated for 1h in ALI-differentiation medium supplemented with 5 µg/ml polybrene (Millipore) before infection.

To generate ACE2-GFP knock-ins in differentiated HNEC, cells were transduced with 20 µl of each concentrated lentivirus, the first produced from pFUGW-spCas9, carrying spCas9 and the second produced from pFUGW-hsACE2-GFP-KI_mCherry-KASH, carrying the sgRNA targeting ACE2 exon 18, the EGFP donor module and the marker mCherry-KASH, labeling infected cells.

### Sample preparation

#### Immunofluorescence staining of fixed samples

Prior to fixation, HNEC were apically washed with PBS. Vero E6 cells and HNEC were fixed in 4% paraformaldehyde and 4% sucrose in PBS (w/v, w/v) for 15 min at 37°C or 20 min with prewarmed fixative for infected samples, permeabilized in 0.5% Triton X-100 in PBS (v/v) for 15 minutes and blocked in 3% BSA in PBS (w/v) for 30 min Before staining of HNEC, filters were cut out from transwell chambers and sectioned into quarters. HNEC on filter fragments were suspended in a 30 µl droplet of primary antibodies diluted in 3% BSA in PBS (w/v) overnight at 4°C. Then, samples were washed three times in 0.1% Triton X-100 in PBS (v/v) for 20 minutes before the addition of corresponding secondary antibodies Alexa Fluor 488-, Alexa Fluor 594, Atto 647N, or Abberior STAR635p-conjugated anti-rabbit and anti-mouse together with DAPI and Phalloidin-Atto-647N (1:200) for 2 hours at room temperature. Cells were washed three times with 0.1% Triton X-100 in PBS (v/v) for 10 minutes. When staining dsRNA, all buffers were supplemented with RNAse inhibitor (RNasin, #N2111, Promega, 1:320.000). Cells were mounted in Prolong Diamond (Thermo Scientific, #P36970), and for airway cultures, covered with precision 13 mm glass coverslips (thickness No. 1.5H, #0117580, Marienfeld).

### TREx protocol, mCLING treatment and general protein stains

For Ten-fold Robust Expansion Microscopy (TREx), cells were either fixed for 20 minutes with pre-warmed (37°C) 4% paraformaldehyde for specific antibody labeling in combination with general protein stains or 4% paraformaldehyde (w/v) and 0.1% glutaraldehyde (w/v) in PBS for visualization of lipid membranes. For mCLING treatment, cells were washed twice in PBS after fixation and incubated in 5 μM mCLING-Atto647N (Synaptic Systems, 710 006AT1) in PBS overnight at room temperature. The following day, cells were fixed a second time with pre-warmed (37°C) 4% paraformaldehyde (w/v) and 0.1% glutaraldehyde (w/v) in PBS. Next, cells were washed with PBS and permeabilized using 0.2% Triton X-100 in PBS (v/v) for at least 15 min. Epitope blocking and antibody labeling steps were performed in 3% BSA in PBS (w/v).

TREx was performed as described earlier [28]. In brief, samples were post-fixed with 0.1 mg/mL acryloyl X-SE (AcX; Thermo Fisher, A20770) in PBS overnight at room temperature. For gelation, monomer solution was prepared containing 1.085 M sodium acrylate (Sigma-Aldrich, 408220), 2.015 M acrylamide (AA; Sigma-Aldrich, A4058) and 0.009% N,N’-methylenebisacrylamide (BIS; Sigma-Aldrich, M1533) in PBS. Gelation of the monomer solution was initiated with 0.15% ammonium persulfate (APS) and 0.15% tetramethylethylenediamine (TEMED). To expand filter-cultured HNECs, 80 μl of gelation solution was pipetted directly in the Transwell chamber set down on a parafilm-covered glass slide. The sample was directly transferred to a 37°C incubator for 1 h to fully polymerize the gel without closing the gelation chamber. All gels, except for samples that were processed for subsequent general protein staining, were transferred to a 12-well plate and digested in TAE buffer (containing 40 mM Tris, 20 mM acetic acid and 1 mM EDTA) supplemented with 0.5% Triton X-100, 0.8 M guanidine-HCl, 7.5 U/mL Proteinase-K (Thermo Fisher, EO0491) and DAPI for 4 h at 37°C. The gel was transferred to a Petri dish, water was exchanged twice after 30 min and the sample was left in water to expand overnight in MiliQ.

For the general protein stains, gels were washed after polymerization in PBS for 15 min at least twice. Next, the gels were incubated for 1 h on a shaker at room temperature with either with 20 μg/mL Atto 594 NHS ester (Sigma-Aldrich, 08471) or 20 μg/mL Atto 647N maleimide (Sigma-Aldrich, 41784) in PBS, both prepared from 20 mg/mL stock solutions in DMSO, to label lysines and cysteines, respectively. These stainings highlighted various subcellular structures to different degrees, which motivated our choice for one or the other in specific experiments. For example, NHS ester strongly labeled basal bodies, whereas maleimide nicely labeled membranous organelles. After staining, gels were washed with an excess of PBS, subsequently digested for 4 h at 37°C as described above, transferred to a Petri dish and expanded overnight. Prior to imaging, the gels were trimmed and mounted.

### Antibodies

Primary antibodies used in this study were rabbit anti-ACE2 (Sino Biological, #80031-RP02), mouse beta-tubulin IV (BioGenex, #MU178-UC), rabbit anti-Calnexin (Abcam, ab22595), mouse anti-CC10 (Origene, AM26360PU-N), Mouse-anti-CD63 (DSHB Hybridoma Product H5C6, deposited to the DSHB by August, J.T. / Hildreth, J.E.K.), Mouse anti-Centriolin antibody (C-9, Santa Cruz Biotechnology, sc-365521), mouse anti-dsRNA (J2, Sci Cons, #10010200), mouse anti-GFP (3E6, Thermo Fisher Scientific, #A-11120), mouse anti-GM130 (BD Biosciences, #610823), mouse anti-MUC5AC (Thermo Fisher Scientific, #MA1-38223), rabbit anti-MUC5AC (Abcam, ab198294), rabbit anti-ODF2 (Abcam, ab43840), rabbit anti-p63 (Abcam, ab124762), mouse anti-SARS CoV/SARS-CoV-2 Nucleocapsid (Sino Biological, #40143-MM08), rabbit anti-SARS CoV Nsp3 (Rockland, #100-401-A52), rabbit anti-SARS-CoV-2 Spike S1 (Sino Biological, # 40150-R007) and rabbit anti-TMPRSS2 (Atlas Antibodies, Cat# HPA035787). Secondary antibodies were highly cross-adsorbed Alexa Fluor -488 or -594-conjugated goat antibodies against rabbit IgG or mouse IgG, IgG1 or IgG2A (Thermo Fisher Scientific), goat-anti-mouse and goat anti-rabbit Abberior STAR 635P and goat anti-mouse Atto 647N (Sigma-Aldrich), together with Atto 647N Phalloidin (ATTO-TEC) and DAPI (Sigma-Aldrich).

### Image acquisition and analysis

#### Confocal and STED imaging of non-expanded samples

Data from non-expanded samples shown in Figures 2C-E, 6C and S1 were acquired using a Zeiss LSM880 Fast Airyscan microscope with 405 nm, 561 nm, 633 nm and argon multiline lasers, internal Zeiss spectral 3 PMT detector and spectroscopic detection using a Plan-Apochromat 63x/1.2 glycerol objective.

Data from non-expanded samples shown in Figures 1, 2F-J, 4A, 5A, 6A, 6B, 6F, 6G and S2 were acquired using a Leica SP8 STED 3X microscope with 405 nm and pulsed (80 MHz) white-light lasers, PMT and HyD detectors and spectroscopic detection using a HC PL APO 93x/1.3 GLYC motCORR STED (Leica 11506417) glycerol objective. For STED imaging of Alexa Fluor 594, Atto674N and Abberior STAR635p, we used 594 nm and 633 nm laser lines for excitation and a 775 nm synchronized pulsed laser for depletion. For Alexa 488 we used 488 nm excitation and a 592 nm continuous depletion laser line. For multicolor STED imaging, each fluorescent channel was imaged using the 2D STED configuration in sequential z-stack mode from highest to lower wavelength to prevent photobleaching. The z-stacks were subjected to mild deconvolution using Huygens Professional software version 17.04 (Scientific Volume Imaging, The Netherlands) with the classic maximum likelihood estimation (CMLE) algorithm and the signal-to-noise ratio (SNR) parameter equal to 7 over 10 iterations.

### Expanded sample imaging

Expanded gels were imaged using the same Leica TCS SP8 STED 3X microscope pulsed (80 MHz) with 405 nm and pulsed (80 MHz) white-light lasers, PMT and HyD detectors and spectroscopic detection with a HC PL APO 86x/1.20W motCORR STED (Leica 15506333) water objective. Images in Figures 5B and S3 were acquired using a Zeiss Lattice Lightsheet LL7 pre-serial system, equipped with a 13.3x/0.44 excitation objective and a 44.83x/1 observation objective. For illumination we used Sinc3 beams with 100 mm length and 1800 nm thickness (without side lobes), dithered in the plane of lightsheet illumination. Images were projected to the pco.edge 4.2 sCMOS camera with a final pixel size of 145 nm. Volumes were acquired with the stage scanning by step size of 0.7 µm parallel to the coverslip.

### Basal Body analysis

Basal body (BB) alignment and polarization were determined using human nasal epithelial cells stained for Odf2 and Centriolin, markers for the BB and basal foot (BF), respectively [70]. For BB alignment, cells were manually grouped into three categories based on the BB pattern of the majority of the cell. Cells with a “Floret” pattern are characterized by small, typically round clusters of regularly spaced BBs without the presence of neat rows. Cells grouped as “Scattered” show random BB localization without clear grouping, often with isolated BBs. “Aligned” cells have short rows of BBs with neat spacing between the BBs in a given row and rows are mostly parallel to each other. The localization of Centriolin puncta relative to the Odf2 signal was used to determine the orientation of BBs. The orientation of the majority of the BBs within one cell was used to categorize cells into one of the three BB polarization groups. Cells with “Random” BB polarization showed no uniform BF position between BBs, even when comparing neighboring BBs. “Locally polarized” cells contain small groups of neighboring BBs that are similarly oriented, but the cell as a whole does not have a single uniform beating direction. Cells are “Polarized” when there is a clear and consistent beating direction such that all BBs are similarly oriented, possibly with the exception of a few isolated BBs.

### TREx data analysis

Processing and analysis of data from expanded samples was done using ImageJ and Arivis Vision4D (Arivis AG) for 3D visualization and rendering. All single planes shown in this manuscript are sum projections of 3 planes (z-spacing 0.36 or 0.15 μm), expect when volumetric rendering is indicated. Contrast was adjusted manually. Figures 3E, 6D and 6E show volumetrically rendered stacks depth-coded in grayscale along the z-axis using Arivis to provide contrast reminiscent of scanning electron microscopy. All volumetric renders except Figure 5 are running sum projections of 3 planes generated using the RunningZProjector plugin (https://valelab4.ucsf.edu/~nstuurman/IJplugins/Running_ZProjector.html) for ImageJ after which the data was imported into Arivis. For Figure 3D gamma was adjusted manually to increase visibility of both cilia and intracellular structures. Some rendered data was filtered in Arivis using the Discrete Gaussian Filter with smoothing radius of 2 to aid visibility.

For Figure 4C and single planes and zooms of 4D and 4E, raw data was imported into ImageJ. Acquisitions were processed using a 3D gaussian blur with a sigma value x, y and z of 0.5 or 2 for the maleimide channel and Spike/GM130/DAPI channels, respectively. Indicated planes are sum projections of 3 planes (z-spacing 0.36 μm) of the filtered stack. For all images, the maleimide channel is shown in inverted contrast. To visualize GM130 and Spike in combination with the inverted maleimide channel, multicolor images of maleimide with GM130 and Spike were presented as overlays instead of merges. In brief, manual intensity-based thresholding of GM130 and Spike channels was used to generate binary masks of these respective channels, which were then converted to regions of interest (ROI). The merged images depict the non-thresholded Spike and GM130 images, merged with the maleimide, but the maleimide signal is not shown within these GM130 and Spike-positive ROIs. For Figure 4 H-I, virus-induced large organelles were defined as discrete bulbous organelles with internal structure that were morphologically distinct from other cellular structures. The diameter corresponds to the largest width of the structure.

Figure 5C is a volumetric render of the raw dataset. For the right panels of Figure 5C, the draw object function in Arivis was used to manually segment the MVB outer membrane, which was rendered as an object (in green) in the far-right plane and used for cropping the dataset in the lower panel to show only the MVBs. The diameter corresponds to the largest width of the structure. To segment individual ILVs, cropped MVBs were first morphologically segmented by the Arivis machine learning algorithm into separate objects (see Figure 5E, right panel, manually drawn ROIs were used for training), objects smaller than 0.1 µm^3^ were filtered out and the remaining objects were inclusion-filled, split using a watershed algorithm and visually inspected to be discrete ILVs and for the presence of Spike. Arivis was used to calculate the bounding box lengths for each ILV.

For segmentation in Figure 7G, the raw dataset was imported in ImageJ and was segmented for microvilli and cilia using the trainable Weka segmentation plugin [71] (https://zenodo.org/record/59290) in ImageJ.

## ACKNOWLEDGEMENTS

We thank Harold MacGillavry and Jelmer Willems for providing plasmids and help with the ORANGE system, and Nalan Liv for the CD63 antibody. This work was supported by the European Research Council (ERC Consolidator Grant 819219 to L.C.K) and the Eindhoven-Wageningen-Utrecht Alliance (www.ewuu.nl) that supports the Centre for Living Technologies. The authors declare no competing financial interests. Additional funding came from the PPP allowance made available by Health∼Holland, Top Sector Life Sciences & Health to stimulated public-private partnerships. The authors declare no competing financial interests.

## SUPPLEMENTAL FIGURES

**Supplemental Figure 1, related to figure 1.**
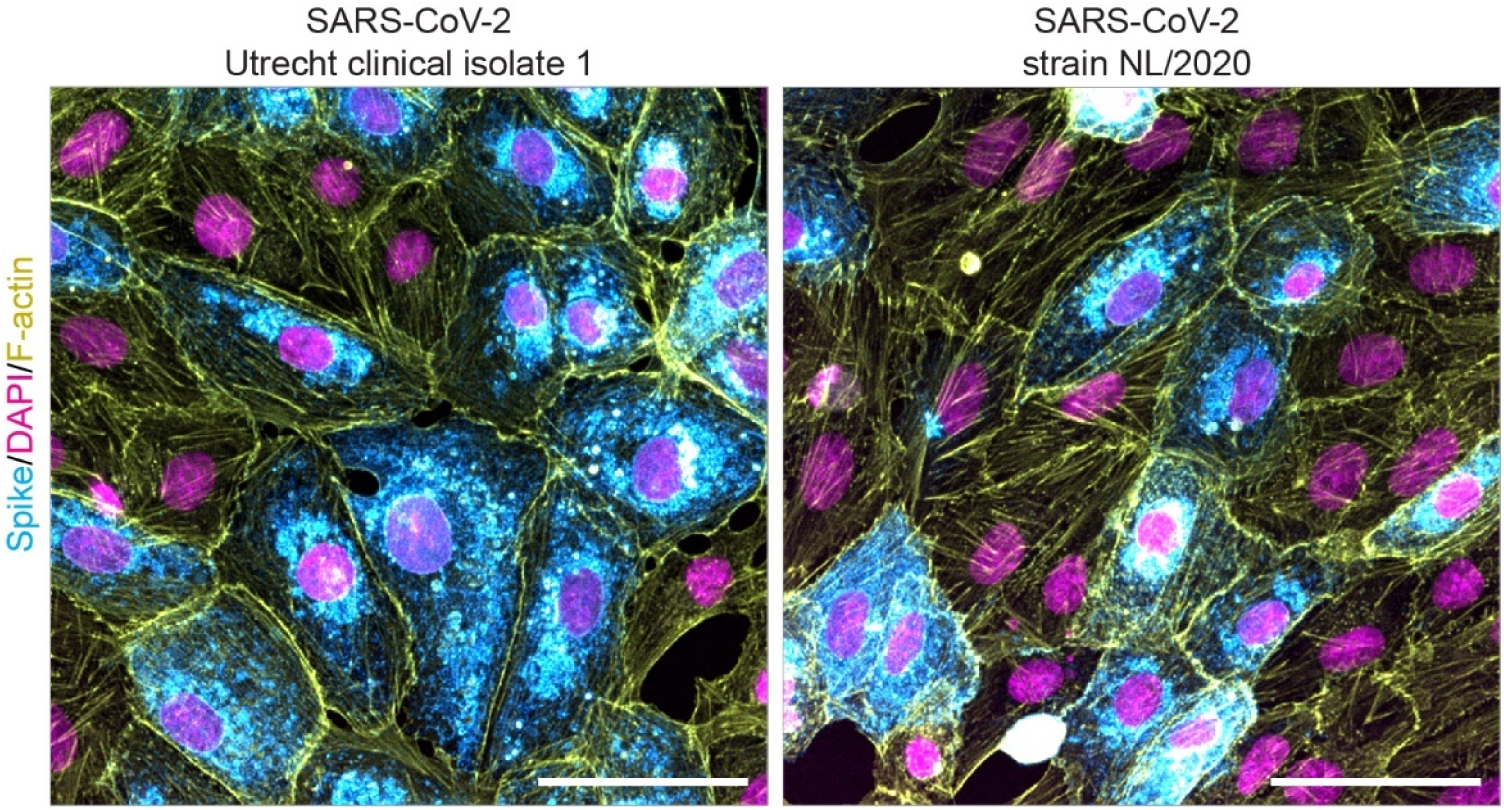
Vero E6 cells infected with different SARS-CoV-2 strains. Immunofluorescent imaging of spike, nuclei (DAPI) and F-actin in Vero E6 cells infected with SARS-CoV-2 strains used in this study. MOI 0.01, cells were fixed at 1 dpi. Scale bars are 50 µm.

**Supplemental Figure 2, related to figure 2.**
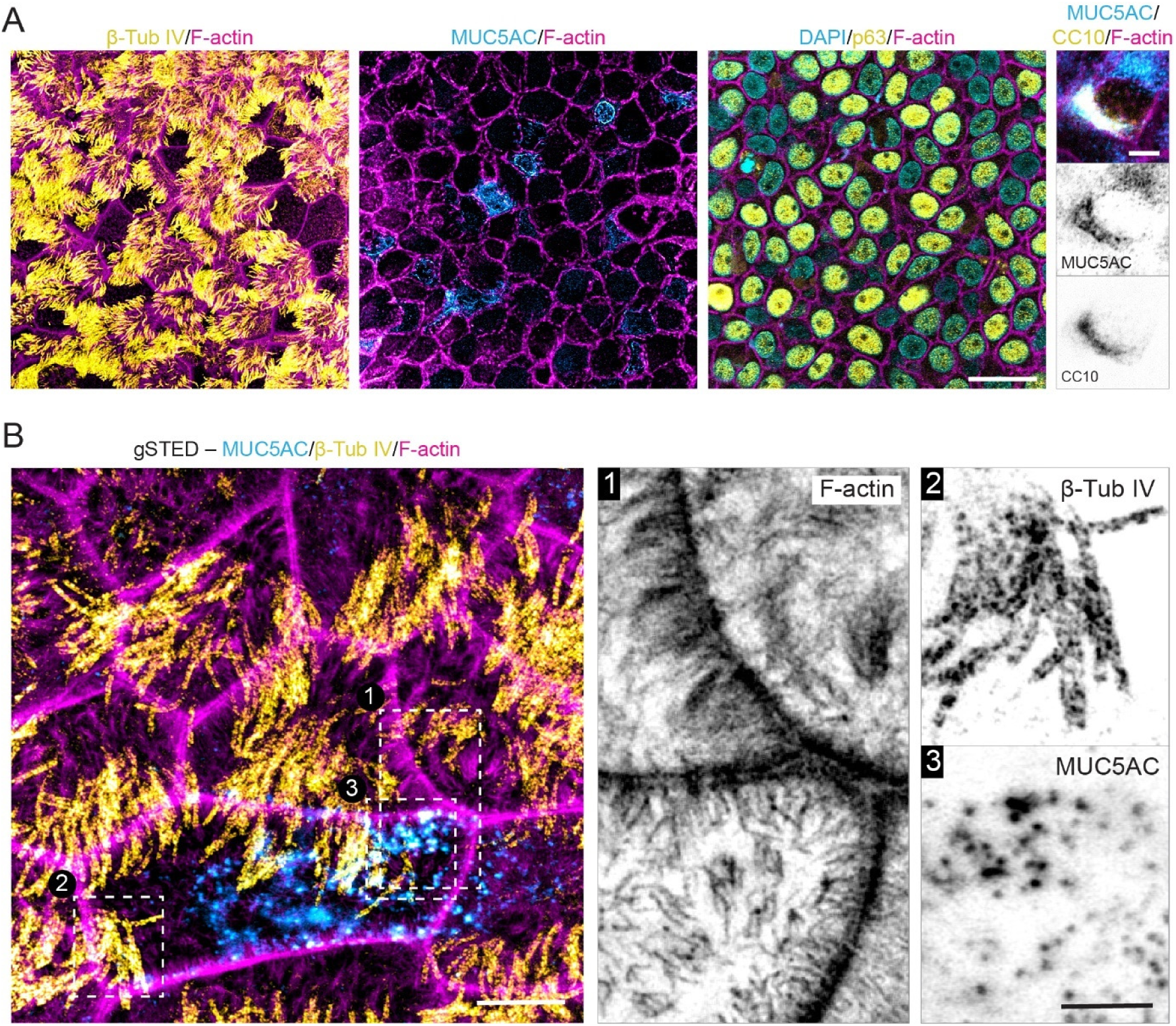
Characterization of human airway cultures. (A) Validation of differentiated airway cultures used in this study by immunofluorescent imaging of markers for distinct cell types: β-Tubulin IV for ciliated cells, MUC5AC for goblet cells, p63 for basal cells and a combination of MUC5AC and CC10 for club cells. (B) gSTED imaging of F-actin (1), B-tub IV (2) and MUC5AC (3) in differentiated HNECs. Scale bars are 20 µm (A overview), 5 µm (A zoom and B overview), or 2 µm (Zoom B).

**Supplemental Figure 3, related to figure 5.**
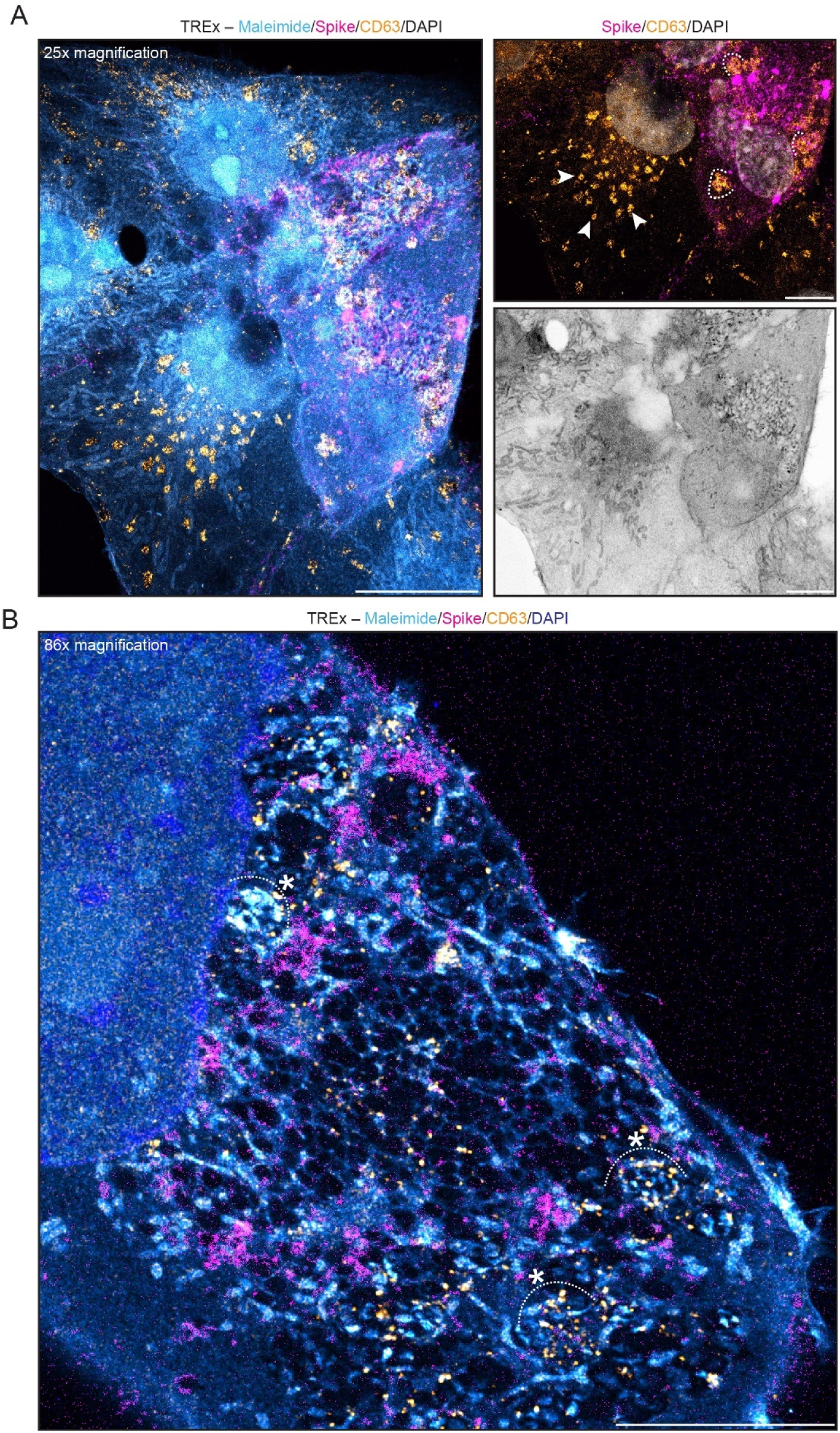
Multi-scale TREx imaging of SARS-CoV-2 infected Vero E6 cells. (A,B) TREx imaging of cellular ultrastructure and indicated markers in SARS-CoV-2 infected and uninfected Vero E6 cells in a large field of view, imaged using a 25x/0.8 water objective (A) or with suborganellar resolution using a 86x/1.20 water objective (B), showing small and large CD63-positive multivesicular bodies in uninfected and infected cells, respectively (arrow heads and curved lines). A detail of this overview is depicted in figure 5B. Scale bars are 10 µm (A), 5 μm (Zooms in A, B). TREx scale bars were corrected to indicate pre-expansion dimensions. MOI: 0.01. Cells were fixed at 2 dpi.

**Supplemental Figure 4, related to figure 7.**
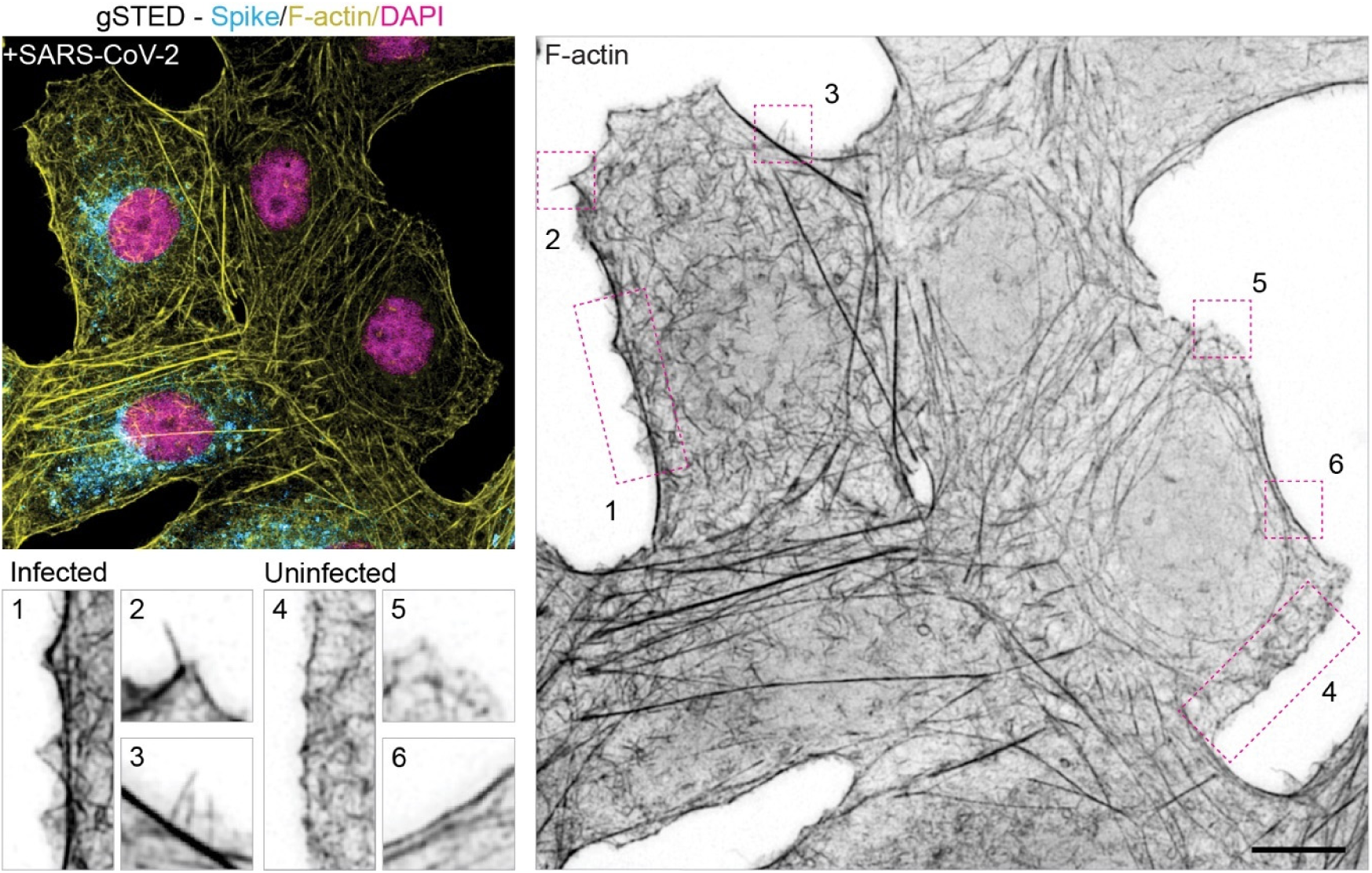
Actin reorganization in infected Vero E6 cells. (A) gSTED imaging of Spike, F-actin (phalloidin) and nuclei (DAPI) in SARS-CoV-2 infected Vero E6 cells. Representative zooms show regions of increased filopodia and membrane ruffling compared to similar regions in non-infected cell.

**Supplemental Figure 5, related to figure 7.**
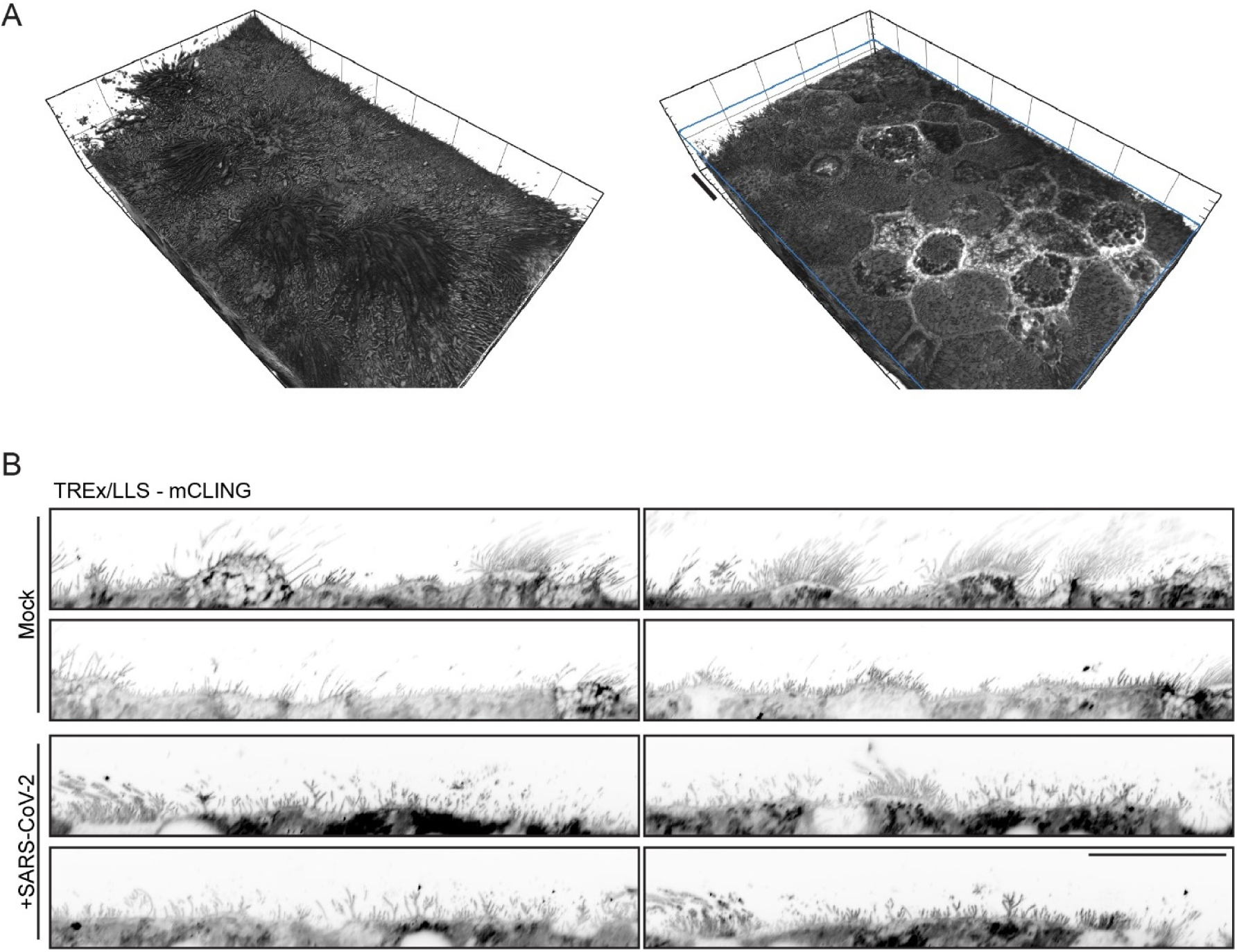
Lattice-light sheet acquisition of mock- and SARS-CoV-2 infected airway cells. (A) Render of airway cultures processed with TREx in combination with mCLING. The complete 2.1mm x 3.5mm area was captured during one acquisition using a lattice-light sheet microscope. (B) Representative side views of mock (top) and SARS-CoV-2 (bottom) infected airway cultures expanded and stained with mCLING. Scale bars are 40 µm (A) 10 µm (B), MOI: 0.2. Cells were fixed at 3 dpi.

## SUPPLEMENTAL VIDEO LEGENDS

**Supplemental movie 1:**

This video corresponds to Figure 3B-E. TREx imaging of human airway cells stained with membrane (blue) and protein (magenta) dyes. Various clipping planes are used and single sections are presented to show the utility of this technique in visualizing complex structures, especially through the use of total membrane and protein stains to provide structural context.

**Supplemental movie 2:**

This video corresponds to Figure 4B. TREx imaging of human airway cells infected with SARS-CoV-2 stained for total proteins (blue), Spike (magenta), and Golgi (yellow). Orange arrowheads mark virus-induced large MVB-like organelles. The dispersion of the Golgi (GM130) is apparent, as well as the close association of Spike with these dispersed structures.

**Supplemental movie 3:**

This video corresponds to Figure 4D. TREx imaging of human airway cells infected with SARS-CoV-2 stained for total proteins (inverted greyscale) and Spike (magenta). Orange arrowheads indicate virus-induced intracellular structures. Circles and arrowheads indicate virus-induced large MVB-like organelles

**Supplemental movie 4:**

This video corresponds to Figure 5C. TREx imaging of Vero E6 cells infected with SARS-CoV-2 stained for total proteins (blue), Spike (magenta) and CD63 (yellow/orange). Large Spike densities are apparent, as well as atypical multivesicular bodies. The external surface of these MVBs is highlighted in green, and these structures are subsequently shown in isolation. An MVB that appears to fuse with the plasma membrane is circled. Here, arrowheads mark intralumenal vesicles visible from the outside of the cell.

**Supplemental movie 5:**

This video corresponds to Figure 7A. TREx imaging of Vero E6 cells infected with SARS-CoV-2 stained for total proteins (greyscale) and Spike (magenta), showing filopodia on Spike-positive cells, and revealing the presence of Spike along these filopodia and on the cell surface. Membrane sheets are also apparent.

**Supplemental movie 6:**

This video corresponds to Figure 7G,H. TREx imaging of human airway cells infected with SARS-CoV-2 stained for total proteins (blue) and Spike (magenta). Orange arrowheads indicate microvilli with Spike puncta along them. The Spike staining (magenta) is then shown superimposed on morphologically segmented microvilli (yellow) and cilia (green), highlighting the overlap of Spike with microvilli and not with cilia.

